# Novel structural components generate distinct type VI secretion system anchoring modes

**DOI:** 10.1101/2020.04.29.069310

**Authors:** Patricia Bernal, R. Christopher D. Furniss, Selina Fecht, Rhoda C.Y. Leung, Livia Spiga, Despoina A.I. Mavridou, Alain Filloux

**Affiliations:** MRC Centre for Molecular Bacteriology and Infection, Department of Life Sciences, Imperial College London, London, SW7 2AZ, UK; Department of Molecular Biosciences, University of Texas at Austin, Austin, 78712, Texas, USA; Department of Biology, Faculty of Sciences, Universidad Autónoma de Madrid, Madrid, 28049, Spain; Departamento de Microbiología, Facultad de Biología, Universidad de Sevilla, Seville,41012, Spain

**Keywords:** Type VI secretion system, interbacterial competition, contact-dependent killing, toxin delivery, TssA, TagB, TagA, sheath stabilization, sheath contraction, *Pseudomonas*

## Abstract

The type VI secretion system (T6SS) is a phage-derived contractile nanomachine primarily involved in interbacterial competition. Its pivotal component, TssA, is indispensable for the assembly of the T6SS sheath structure, the contraction of which propels a payload of effector proteins into neighboring cells. Despite their key function, TssA proteins exhibit unexpected diversity and exist in two major forms, a short (TssA_S_) and a long (TssA_L_) TssA. Whilst TssA_L_ proteins interact with a partner, called TagA, to anchor the distal end of the extended sheath, the mechanism for the stabilization of TssA_S_-containing T6SSs remains unknown. Here we discover a novel class of structural components that interact with short TssA proteins and contribute to T6SS assembly by stabilizing the polymerizing sheath from the baseplate. We demonstrate that the presence of these components is important for full sheath extension and optimal firing. Moreover, we show that the pairing of each form of TssA with a different class of sheath stabilization proteins results in T6SS apparatuses that either reside in the cell for a while or fire immediately after sheath extension, thus giving rise to different aggression behaviors. We propose that this functional diversity could contribute to the specialization of the T6SS to suit bacterial lifestyles in diverse environmental niches.

## INTRODUCTION

Bacteria live in complex polymicrobial communities that are shaped by interspecies cooperation and competition. As resources are limited, antagonistic strategies are a major driver of survival and success for bacterial populations. One of the most elaborate bacterial weapons is the type VI secretion system (T6SS), which not only promotes inter-bacterial and inter-kingdom competition (1–3), but is also involved in the interaction of bacteria with their hosts (4, 5) and the acquisition of both nutrients (6, 7) and genetic material (8, 9).

The T6SS apparatus is a contractile nanomachine that delivers proteinaceous effectors to neighboring cells in a contact-dependent manner (10, 11). The system is a multiprotein complex (12), which when fully assembled, extends across the entire width of the cell (13) (Fig. 1A). Three main components make up this structure: the membrane complex, the baseplate and the tail, comprising the contractile sheath and the inner tube. The membrane complex (TssJLM) spans the cell envelope (14–16), providing a platform on to which the baseplate (TssEFGK) docks (17, 18). After the baseplate is in position it promotes the polymerization of the contractile sheath (TssBC), which encompasses the Hcp tube topped by the VgrG/PAAR complex (19–21). Upon sheath contraction (22) the Hcp tube and the VgrG tip, along with their associated effectors, are propelled outside the attacker and into neighboring cells (10, 11). This is followed by ClpV-dependent sheath disassembly and recycling before another round of firing (23–25) (Fig. 1A).

**Fig. 1.**
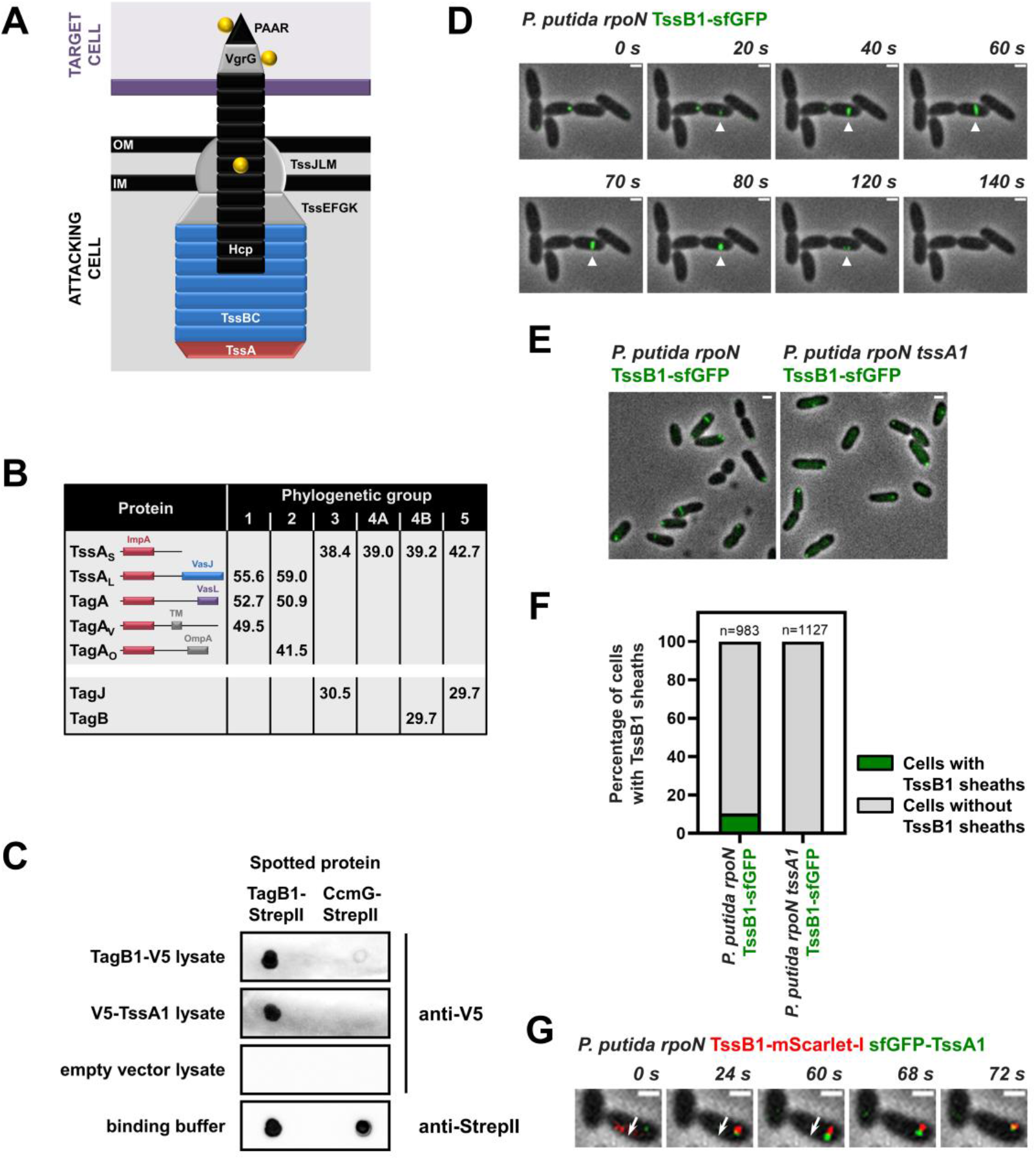
Short TssA proteins interact with novel T6SS structural components. **(A)** Schematic representation of a T6SS apparatus at the moment of firing. The Hcp tube (in black) along with the VgrG/PAAR tip complex (in grey/black) and their associated effectors (in yellow) are propelled from the attacking cell into a neighboring target cell (in purple). The membrane complex and baseplate structures are shown in grey, whilst the sheath and the cap protein TssA are depicted in blue and red, respectively. **(B)** Synopsis of the results from the *in silico* analysis presented in File S1. Average protein sizes in kDa are given for T6SS components of each phylogenetic group and the domain architecture of TssA-like proteins is shown; the N-terminus of all TssA-like proteins forms an ImpA domain whilst their C-terminus varies (28, 29). “TagA” refers to homologues of the *E. coli* TagA anchor protein (30), “TagA_V_” has been described as the anchor for the T6SS of *V. cholerae* (29), and “TagA_O_”, which has a different C-terminus compared to the other two forms of TagA, has not been described before. Long TssA proteins (TssA_L_) are encoded in T6SS clusters from phylogenetic groups 1 and 2, whilst short TssA proteins (TssA_S_) are encoded in T6SS clusters from phylogenetic groups 3-5. Representative examples of well-studied T6SSs from each phylogenetic group are: *P. aeruginosa* H2-T6SS and *V. cholerae* T6SS (group 1); EAEC T6SS (group 2); *P. aeruginosa* H1-T6SS (group 3); *P. aeruginosa* H3-T6SS (group 4A); *P. putida* K1-T6SS (group 4B); *Agrobacterium tumefaciens* T6SS (group 5). TssA_L_ proteins usually co-occur with TagA-like components (only one TagA protein and its cognate TssA partner are encoded in each cluster), but examples of T6SS clusters belonging to phylogenetic groups 1 and 2 encoding a TssA_L_ without containing a *tagA* gene also exist (for example the H2-T6SS of *P. aeruginosa* in phylogenetic group 1 (29)). TssA_S_ proteins often co-occur with T6SS components that are smaller in size and are largely uncharacterized (TagJ and TagB proteins). As with TssA_L_, there are clusters where no gene encoding a TssA_S_ partner protein is identified (for example all clusters from phylogenetic group 4A or some members of phylogenetic group 5). **(C)** Far-western dot blot assays showing interaction of *P. putida* TagB1 with itself and its cognate TssA1. Purified TagB1-StrepII was spotted on a nitrocellulose membrane and exposed to *E. coli* cell lysates expressing TagB1-V5 or V5-TssA1. The TagB1 interactions are specific as no interactions with the tested lysates are detected when the binding control protein CcmG-StrepII is probed or when TagB1-StrepII is exposed to lysates from cells harboring the empty vector; similar amounts of TagB1-StrepII and CcmG-StrepII were spotted on the membrane. Levels of StrepII-tagged proteins were assessed using a Strep-Tactin-HRP conjugate and levels of V5-tagged proteins were detected with an anti-V5 antibody and an HRP-conjugated secondary antibody. **(D-G)** *P. putida* **K1-T6SS sheaths display characteristic T6SS behaviors. (D)** *In vivo* imaging of *P. putida rpoN* TssB1-sfGFP. The full cycle of T6SS assembly, including sheath initiation, extension, contraction and disassembly, over approximately 120 s, is shown for a representative sheath (white arrowhead). The panels presented are selected images from a fluorescence microscopy time-lapse recording of *P. putida rpoN* expressing TssB1-sfGFP from the native *tssB1* locus (see Video S1). Images were recorded every 2 s and scale bars represent 1 μm. **(E)** Formation of *P. putida* sheaths is dependent on TssA1. Representative fluorescence microscopy images of *P. putida rpoN* and *P. putida rpoN tssA1* expressing TssB1-sfGFP from the native *tssB1* locus. Scale bars represent 1 μm. **(F)** Quantification of (E). Approximately 10% of cells form sheaths in *P. putida rpoN*, whilst no sheaths are detected in the isogenic *tssA1* mutant. n indicates the number of cells included in the analysis. **(G)** *P. putida* TssA1 localizes to the site of sheath initiation and subsequently migrates at the end of the polymerizing sheath. The white arrow indicates the direction of sheath extension. The panels presented are selected images from a fluorescence microscopy time-lapse recording of *P. putida rpoN* expressing sfGFP-TssA1 and TssB1-mScarlet-I from the native gene loci. Scale bars represent 1 μm. Data collection and image analysis protocols for all panels presenting the results of fluorescence microscopy experiments are described in detail in the Materials and Methods.

TssA is a core structural component that is essential for T6SS function (26, 27). It promotes priming and polymerization of the sheath, whereby after recruiting the remaining baseplate proteins, TssA is displaced by the growing sheath and migrates at its distal end, eventually reaching the other side of the cell (27). Because of its central role, TssA is found in all T6SSs, nonetheless it is one of the least conserved core components of the apparatus (File S1; percentile identity of TssA proteins can be low even within the same phylogenetic group). Notably, TssA exists in two major forms, a long 55-60 kDa protein (TssA_L_) and a shorter approximately 40 kDa version (TssA_S_) (28, 29) (Fig. 1B). The structure and function of prototypical TssA_L_ proteins have been extensively studied (27–29) and it is known that the *Escherichia coli*, the *Vibrio cholerae* and the *Aeromonas hydrophila* TssA_L_ form a ring-like structure, where the C-terminus of the protein largely occupies the center of the ring (27–29). It has been recently shown that these long TssA forms interact with an accessory TssA-like protein, called TagA, which secures the distal end of the sheath at the opposing cell membrane, ensuring optimal sheath polymerization and maintenance of the sheath in its extended conformation for long periods of time (29, 30). In contrast, *Pseudomonas aeruginosa* and *Burkholderia cenocepacia* TssA_S_, whilst still ring-like, display structures where the center of the ring is empty (26, 28). In this case, TagA is not present and it is unknown how the sheath is stabilized and anchored. In order to understand the implications of TssA diversity for T6SS assembly dynamics and function it is necessary to elucidate the relationship between short TssA proteins, the baseplate and the extending sheath, and to determine how TssA_S_-containing T6SSs are stabilized.

## RESULTS

### Short TssA proteins interact with novel T6SS structural components

To assess the core composition of T6SSs across different bacterial species, we performed an *in silico* analysis of 100 T6SS clusters (File S1). We found that TssA_L_ proteins dominate phylogenetic groups 1 and 2 and usually co-occur with TagA-like components (Fig. 1B). On the other hand, short TssA proteins, present in phylogenetic groups 3-5, do not co-occur with TagA-like partners, but instead are often found to co-exist with 30 kDa proteins of unknown function (Fig. 1B). This consistent co-occurrence and the proximity of the genes encoding these uncharacterized proteins to the core T6SS genes prompted us to probe their function. We chose *Pseudomonas putida* as a model organism as we have previously shown that in this bacterium a representative of these uncharacterized proteins is encoded by the first gene of the K1-T6SS cluster (31) (Fig. S1), a member of the phylogenetic group 4B also encoding a short TssA protein (Fig. 1B, File S1). The gene of interest (PP_5562 or PP3100.1), previously referred to as *tagX1* (31), will be hereafter called *tagB1* (for type VI accessory gene B1).

TagB1 is a predicted cytoplasmic protein of unknown structure. To identify its T6SS interaction partners we introduced a twin StrepII tag at the C-terminus of the native copy of TagB1 and purified the protein in conditions where the K1-T6SS is active by using an *rpoN* mutant of *P. putida* KT2440 with increased K1-T6SS expression (31). We identified and quantified the eluted proteins by mass spectrometry and found that TssA1 was the only protein significantly enriched in samples containing TagB1-(StrepII)_2_, compared to untagged control samples (File S2). We validated our mass spectrometry results, obtained from the native T6SS apparatus expressed in *P. putida*, by far-western dot blotting. We observed that pure TagB1-StrepII interacted with TssA1-V5 from crude cell extracts and at the same time we detected a clear interaction of TagB1 with itself (Fig. 1C). These associations are specific to TagB1 and TssA1 as no interactions were detected when using a binding control protein or lysates from cells harboring the empty vector (Fig. 1C). The fact that TagB1 self-associates and thus is likely functional in a multimeric state, whilst also interacting with the core component TssA1, suggests that it is a structural T6SS protein.

### P. putida K1-T6SS sheaths display characteristic T6SS behaviors

In order to interrogate whether TagB1 plays a role in the assembly of the K1-T6SS, we visualized the apparatus *in vivo*. The K1-T6SS has been previously shown to be active (31), but it has never been imaged by fluorescence microscopy. We generated a *P. putida rpoN* strain that expresses TssB1 fused to superfolder GFP (sfGFP) from the native *tssB1* locus. Using this strain, we detected sheaths in approximately 10% of cells at any given time (Fig. 1F), and we could visualize the full cycle of T6SS assembly, including sheath initiation, full extension, contraction and disassembly over approximately 120 seconds (Fig. 1D, Video S1). Sheath structures with identical dynamics could also be seen in wild-type *P. putida* expressing TssB1-sfGFP (Fig. S2A), although they occurred at lower frequency.

No sheaths were detected in the absence of TssA1 (Fig. 1EF) consistent with the fact that *P. putida tssA1* has been reported to be essential for K1-T6SS function (31) and that in other organisms sheath formation is only observed *in vivo* when TssA is present (27). As it has been demonstrated that long TssA proteins from *E. coli*, *V. cholerae* and *P. aeruginosa* (H2-T6SS) initially localize at the baseplate and subsequently migrate at the tip of the growing sheath (27, 29), we wanted to investigate if a short TssA protein, like *P. putida* TssA1, exhibits a similar behavior. To monitor its localization during sheath assembly, we generated a *P. putida rpoN* strain expressing sfGFP-TssA1 and TssB1-mScarlet-I from their native loci. TssA1 localized to the site of sheath initiation, before the sheath structure could be detected, and was then displaced to the distal end of the polymerizing sheath, reaching the other side of the cell prior to sheath contraction (Fig. 1G). This is the first time this has been demonstrated for a short TssA protein and suggests that, despite its two distinct forms, the localization of TssA during T6SS assembly and its interplay with the baseplate and sheath proteins is likely conserved.

### TagB1 stabilizes sheath polymerization from the baseplate

Having confirmed that *P. putida* K1-T6SS sheaths display characteristic behaviors (Fig. 1DEFG), we proceeded to investigate the behavior of TagB1 *in vivo*. We generated a *P. putida rpoN* strain that expresses TagB1 fused to sfGFP from the native *tagB1* locus and found that TagB1-sfGFP forms transient foci that accumulate over time before abruptly disappearing (Fig. 2A, Video S2). Similar to sheaths (Fig. 1F), approximately 10% of *P. putida rpoN* cells contain TagB1-sfGFP foci at any given time (Fig. 2C). The same transient foci were also detected in wild-type *P. putida* expressing TagB1-sfGFP (Fig. S2B), but as expected, the percentage of cells exhibiting foci was markedly lower. Importantly, no TagB1-sfGFP foci were observed in the absence of TssA1 (Fig. 2B and 2C, left graph), whilst the number of cells exhibiting TssA1-sfGFP foci remained unchanged in a *tagB1* mutant (Fig. 2C, right graph). This indicates that interaction of TagB1 with TssA1 is necessary for TagB1 activity, whereas the presence of TagB1 is not required for the function of TssA1.

**Fig. 2.**
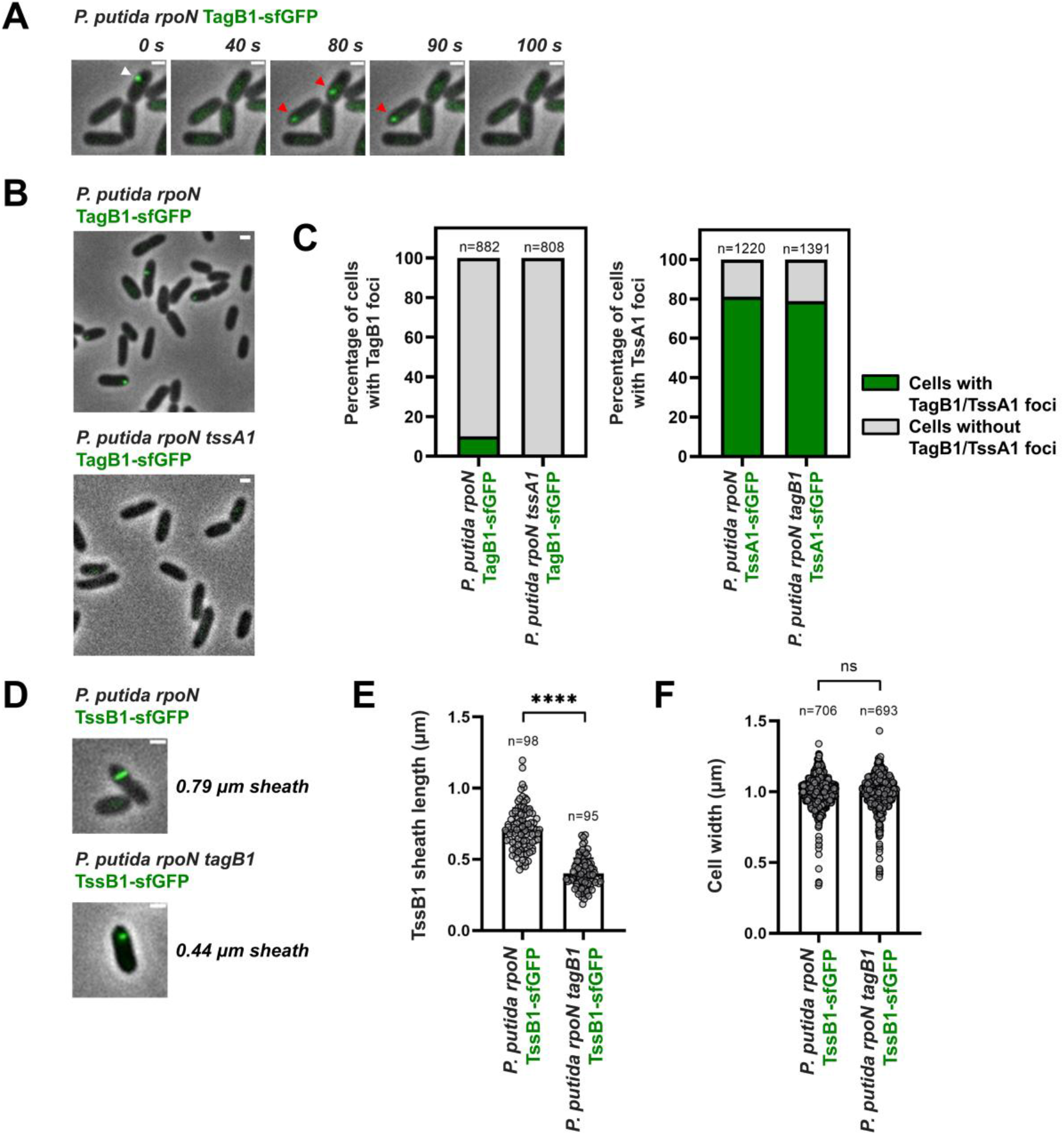
*P. putida* TagB1 is required for the full extension of K1-T6SS sheaths. **(A)** *In vivo* imaging of *P. putida rpoN* TagB1-sfGFP. TagB1 forms transient foci that accumulate over time before abruptly disappearing (white arrowhead). Disappearance of foci is not due to photobleaching as new TagB1-sfGFP foci appear in the same or neighboring cells (red arrowheads). The panels presented are selected images from a fluorescence microscopy time-lapse recording of *P. putida rpoN* expressing TagB1-sfGFP from the native *tagB1* locus (see Video S2). Images were recorded every 2 s and scale bars represent 1 μm. **(B)** Formation of *P. putida* TagB1-sfGFP foci is dependent on TssA1. Representative fluorescence microscopy images of *P. putida rpoN* and *P. putida rpoN tssA1* expressing TagB1-sfGFP from the native *tagB1* locus. Scale bars represent 1 μm. **(C)** The left graph shows quantification of (B); approximately 10% of cells form TagB1-sfGFP foci in *P. putida rpoN*, whilst no foci are present in the isogenic *tssA1* mutant. In contrast to TagB1-sfGFP focus formation, which is dependent on the presence of TssA1, the number of cells containing TssA1-sfGFP foci remains unchanged upon deletion of *tagB1* (right graph). n indicates the number of cells included in the analysis. **(D)** Absence of TagB1 prevents full extension of the K1-T6SS sheath. Representative fluorescence microscopy images of *P. putida rpoN* and *P. putida rpoN tagB1* expressing TssB1-sfGFP from the native *tssB1* locus. Scale bars represent 1 μm. Examples of selected images from fluorescence microscopy time-lapse recordings of *P. putida rpoN tagB1* expressing TssB1-sfGFP are shown in Fig. S3. **(E)** Quantification of (D). The average sheath length in a *tagB1* mutant is approximately 50% smaller than in *P. putida rpoN*. Sheath length measurements were performed manually on fluorescence microscopy time-lapse recordings of *P. putida rpoN* strains expressing TssB1-sfGFP from the native *tssB1* locus; measurements were performed on the frame directly prior to that in which sheath contraction was observed. n indicates the number of cells included in the analysis. **(F)** The cell width of a *tagB1* mutant is identical to that of the parental *P. putida rpoN* strain. Cell width data were directly extracted from manually curated cell masks generated using MicrobeJ. n indicates the number of cells included in the analysis. Data collection and image analysis protocols for all panels presenting the results of fluorescence microscopy experiments are described in detail in the Materials and Methods.

TssA is known to promote T6SS assembly by initiating and assisting sheath polymerization (27). As *P. putida* TagB1 interacts with TssA1 (Fig. 1C, File S2) and TagB1-sfGFP foci are dependent on TssA1 for their formation (Fig. 2BC), we hypothesized that TagB1 could be important for K1-T6SS sheath formation. We imaged sheath dynamics in a *tagB1* mutant expressing TssB1-sfGFP and found that in the absence of *tagB1* sheath length was significantly reduced (Fig. 2DE and S3), with sheaths produced by the *tagB1* mutant only reaching approximately 50% of the length of those produced by the parental strain (Fig. 2E). Since cell width has been shown to control T6SS sheath length (32), we determined the average cell width in the *tagB1* mutant and found it to be identical to that of the parental strain (Fig. 2F). The fact that deletion of *tagB1* affects the maximum length of the sheath points to TagB1 having a role in sheath stabilization, similar to TagA in *E. coli*. Indeed, absence of the TagA anchor also affects the maximum length of the sheath, in this case leading to overly large sheaths that tend to break (30). Therefore, our results suggest that the K1-T6SS, which contains a short TssA, is stabilized through a small (~30 kDa), novel structural component rather than a large TssA-like anchor protein, as is the case for long TssA proteins.

TagB1 interacts with TssA1 (Fig. 1C, File S2), thus it is plausible that it could stabilize the T6SS apparatus from either end of the sheath, since TssA1 localizes initially at the baseplate of the system before being displaced to the other side of the cell via sheath extension. Bacterial-two-hybrid analysis showed that TagB1 interacts with all but one of the baseplate proteins (TssK1, TssE1, TssF1, Hcp1, VgrG1 and TssA1; Fig. 3A). In agreement with this, when we simultaneously visualized TagB1 and TssB1 *in vivo*, TagB1-sfGFP foci were associated with the end of the extended sheath towards which contraction occurred (Fig. S4A). This suggests that TagB1 localizes at the baseplate. Further analysis of the full sheath extension and contraction cycle confirmed this. TagB1-sfGFP foci appeared and accumulated at the site of sheath initiation, after sheath polymerization had started (Fig. 3B and S4B; sheath extension: 0 s - 56 s, TagB1-sfGFP focus appearance: 10 s) and the disappearance of TagB1-sfGFP foci coincided with sheath contraction (Fig. 3B and S4B; sheath contraction: 58 s - 70 s). Together these results (Fig. 3AB and S4) show that TagB1, unlike TagA, localizes at the baseplate and not at the distal end of the sheath. This conclusion is further supported by the fact that after a contraction event had occurred TagB1-sfGFP foci occasionally reappeared in the same position in the cell (Fig. S5). This is much more likely to happen if TagB1 associates with the baseplate rather than the distal end of the sheath and is consistent with the re-use of T6SS membrane complex structures observed by Zoued *et. al.* (27).

**Fig. 3.**
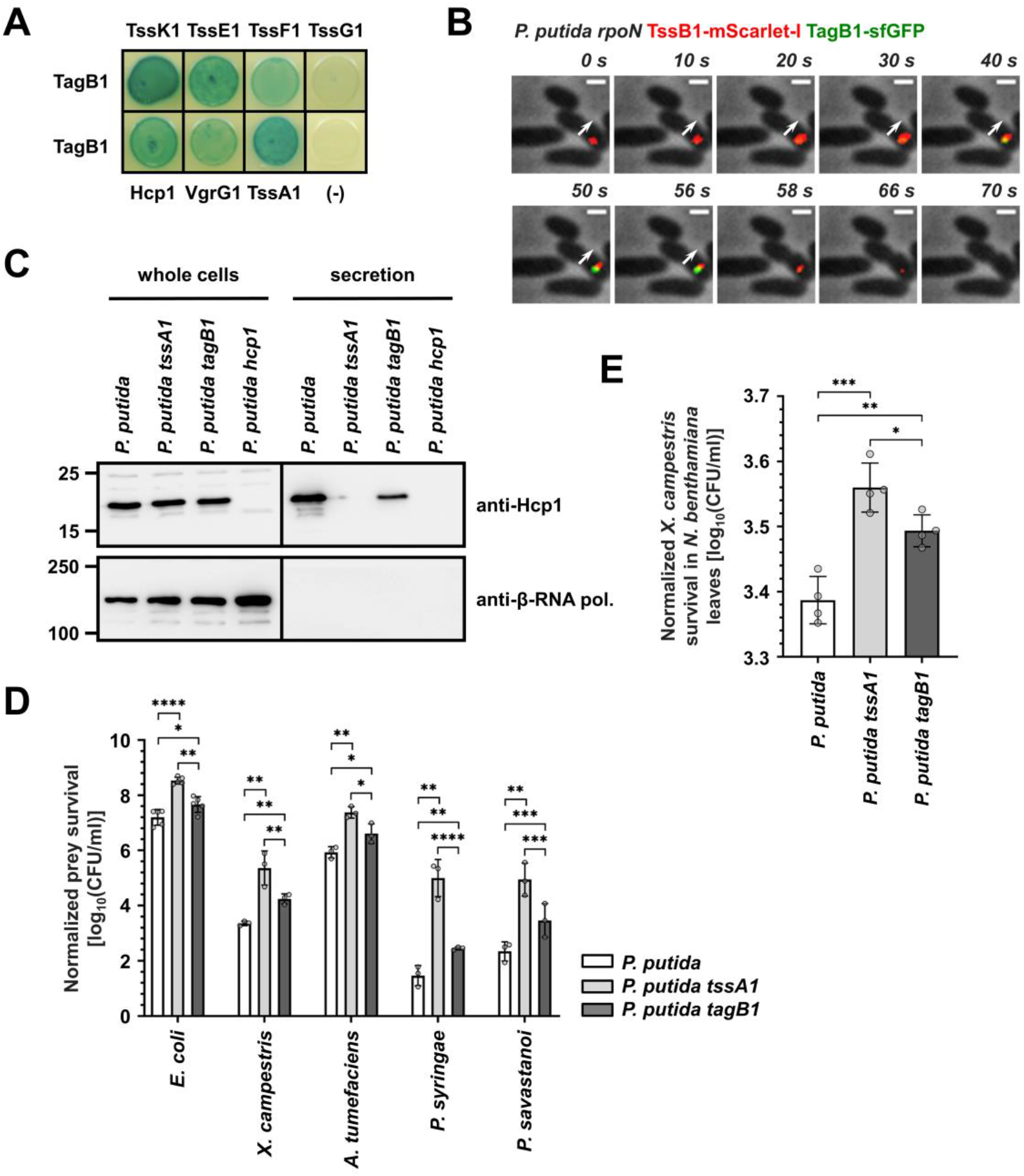
*P. putida* TagB1 anchors the K1-T6SS sheath from the baseplate enabling optimized firing. **(A)** *P. putida* TagB1 interacts with all but one of the baseplate proteins, as assessed by bacterial-two-hybrid assay. BTH101-reporter cells producing the indicated proteins fused to the T18 and T25 domain of the *Bordetella pertussis* adenylate cyclase were spotted on X-Gal-IPTG reporter LB agar plates. Three independent experiments were performed with identical results. **(B)** *P. putida* TagB1 localizes to the baseplate. The white arrow indicates the direction of sheath extension. The panels presented are selected images from a fluorescence microscopy time-lapse recording of *P. putida rpoN* expressing TagB1-sfGFP from the native *tagB1* locus and the gene encoding TssB1-mScarlet-I integrated into the chromosome using the miniCTX plasmid (39) whilst the native *tssB1* copy is still present. Scale bars represent 1 μm. Images from individual fluorescence channels are shown in Fig. S4B. Data collection and image analysis protocols are described in detail in the Materials and Methods. **(C)** A *tagB1* mutant of *P. putida* shows markedly reduced secretion of Hcp1. Hcp1 expression and presence in the culture supernatant were assessed for wild-type *P. putida* and the isogenic *tssA1*, *tagB1* and *hcp1* mutants. Hcp1 protein levels were detected using an anti-Hcp1 primary antibody and an HRP-conjugated secondary antibody. The *P. putida hcp1* mutant was used as a negative control for Hcp1 detection. Positions of molecular weight markers are shown on the left, the β-subunit of the *E. coli* RNA polymerase (β-RNA pol.) was used as a loading and bacterial lysis control and black lines indicate where the membrane was cut. A representative blot from three independent experiments is presented. **(D)** Bactericidal activity of *P. putida* strains against *E. coli* and a panel of plant pathogens. *E. coli, X. campestris*, *A. tumefaciens*, *Pseudomonas syringae* and *Pseudomonas savastanoi* strains harbor the pRL662-gfp plasmid that confers gentamycin resistance. The wild-type *P. putida* strain and the isogenic *tssA1* and *tagB1* mutants were co-incubated with the different prey strains and the outcome of the competitions was quantified by counting colony forming units (CFUs) using antibiotic selection of the input (time = 0 hours) and output (time = 5 hours for *E. coli* or time = 24 hours for plant pathogens). n ≥ 3, graph shows means ±SD, significance is indicated by * = p < 0.05, ** = p <0.01, *** = p < 0.001 or **** = p <0.0001. **(E)** *In planta* competition assay between *P. putida* strains and the phytopathogen *X. campestris*. *X. campestris* (pRL662-gfp) was co-infiltrated with *P. putida* strains on *N. benthamiana* leaves for 24 hours. The outcome of the competition was quantified by counting CFUs using gentamycin selection of *X. campestris* recovered from *N. benthamiana* leaves at time = 0 hours (input) and time = 24 hours (output). n = 4, graph shows means ±SD, significance is indicated by * = p < 0.05, ** = p <0.01 or *** = p < 0.001.

Taken together our data reveal a novel mode of stabilization for this TssA_S_-containing T6SS. In brief, we show that TagB1 is a structural T6SS protein that stabilizes sheath polymerization from the baseplate (Fig. 3AB) and enables the sheath to reach its full size (Fig. 2DEF). For this to happen TagB1 is recruited to the baseplate by TssA1 (File S2, Fig. 1C and 2BC), where it continues to accumulate (Fig. 3AB and S4B, Video S2) through self-interaction (Fig. 1C), supporting the polymerizing sheath after TssA1 has left the site of sheath initiation for the end of the extending structure (Fig. 1G).

### TagB1-mediated sheath stabilization allows optimal K1-T6SS firing

Having determined that TagB1 plays a stabilizing role in the assembly of the T6SS sheath, we wanted to investigate the effect of its absence on K1-T6SS function and during interbacterial competition. We found that compared to a wild-type strain, a *tagB1* mutant exhibited substantially decreased Hcp1 secretion (Fig. 3C). Consistent with this, the *tagB1* mutant was less able to kill both prey *E. coli* and a range of plant pathogens, which *P. putida* would encounter and compete with in its natural environments (31) (Fig. 3D). We observed the same effect during *in planta* competition of *P. putida* strains with the plant pathogen *Xanthomonas campestris* (performed in *Nicotiana benthamiana* leaves) (Fig. 3E). For all assays, the *tagB1* phenotype was not as drastic as that of a *tssA1* mutant, the latter resulting in an entirely inactive T6SS (31) (Fig. 3CDE). This is in agreement with the ability of the *tagB1* mutant strain to form sheaths, which although shorter (Fig. 2DE) are likely to still deliver some effector cargo. Thus, whilst TagB1 is not essential for T6SS activity, it is still a key component of the system as, through its stabilizing role, it mediates optimal firing of the K1-T6SS and effective killing of prey bacteria.

### TagB and TagJ components produce a distinct anchoring mode for T6SSs with short TssA proteins

We have demonstrated that the *P. putida* K1-T6SS is stabilized and achieves optimal function through the recruitment of the small protein TagB1 to the baseplate by TssA1 (a TssA_S_ protein). To determine if this mode of sheath stabilization applies to other T6SSs encoding short TssA components, we selected the H1-T6SS from *P. aeruginosa*, which belongs to phylogenetic group 3 and contains the putative accessory protein TagJ1 (Fig. 1B, File S1). We performed far-western dot blotting with pure TagJ1-StrepII to assess both self interaction and interaction with its cognate TssA_S_ protein. Like *P. putida* TagB1, TagJ1 self interacts and binds strongly to TssA1 (Fig. S6A). In addition, TagJ1-sfGFP forms transient foci *in vivo* (Fig. S6B) which, in a similar manner to *P. putida* TagB1-sfGFP foci, remain static before abruptly disappearing. Therefore, whilst TagJ1 and TagB1 are not homologous, our data indicate that they likely perform similar functions during T6SS assembly. As such, the sheath-stabilization role we described for *P. putida* TagB1 likely extends to other T6SSs encoding short TssA proteins together with TagB, TagJ or other similarly sized partners (Fig. 1B). Notably, this mechanism of T6SS stabilization via baseplate-associated 30 kDa components is completely distinct from the previously reported TagA-mediated anchoring (29, 30) that secures the sheath through clamping of its distal end.

### Distinct anchoring modes generate functionally diverse T6SSs

*E. coli* TagA, which associates with a TssA_L_ protein, has been shown to maintain the assembled sheath in the extended conformation for long periods of time (30, 33). Moreover, the TssA_L_-containing H2-T6SS from *P. aeruginosa* has also been reported to be stable in its extended conformation (29). This contrasts with T6SSs that contain short TssA proteins, like the *P. aeruginosa* H1-T6SS, which is notoriously rapid (29, 34) as well as the *P. putida* K1-T6SS which we find to fire immediately after sheath extension (Video S1). Knowing that the aforementioned T6SS apparatuses have different anchoring mechanisms (TssA_L_-TagA or TssA_S_-TagB/J), we wanted to assess whether there is a correlation between the mode of T6SS stabilization, determined primarily by the type of TssA (TssA_L_ or TssA_S_) and the nature of its associated stabilizing protein, and the time that elapses from the initiation of sheath polymerization until its contraction. To do this, we systematically measured the time to contraction of T6SSs from different phylogenetic groups (*P. aeruginosa* H1-T6SS, group 3; *P. putida* K1-T6SS; group 4B; *P. aeruginosa* H2-T6SS, group 1) (Fig. 4A) and integrated our data with values from the literature (*E. coli* T6SS, group 2 (30); *V. cholerae* T6SS, group 1 (29)) (Fig. 4B). We show that T6SSs with TssA_S_ proteins and a TagB or TagJ partner contract immediately after sheath extension, taking between 20-70 seconds to assemble and fire, whereas systems with TssA_L_ proteins tend to reside in the cell and can remain extended for more than ten minutes (Fig. 4B). Notably, whilst the stability of TssA_L_-containing T6SSs is usually supported by TagA anchors (29, 30, 33), it has been postulated that TssA_L_ components could act as an anchor themselves, for example in the case of the *P. aeruginosa* H2-T6SS (29) for which no TagA protein has been identified to date (File S1). This suggestion is supported by our results, which show that *P. aeruginosa* H2-T6SS sheaths are stably anchored (Fig. 4A, bottom row). Overall, we find that the type of TssA, TssA_L_ or TssA_S_, which in turn drives the recruitment of its respective sheath stabilizing protein partner (TagA or TagB/J), determines the firing dynamics of each T6SS apparatus. As such, the specific pairing of these two structural components likely underpins the different aggression strategies observed across T6SS-carrying bacteria.

**Fig. 4.**
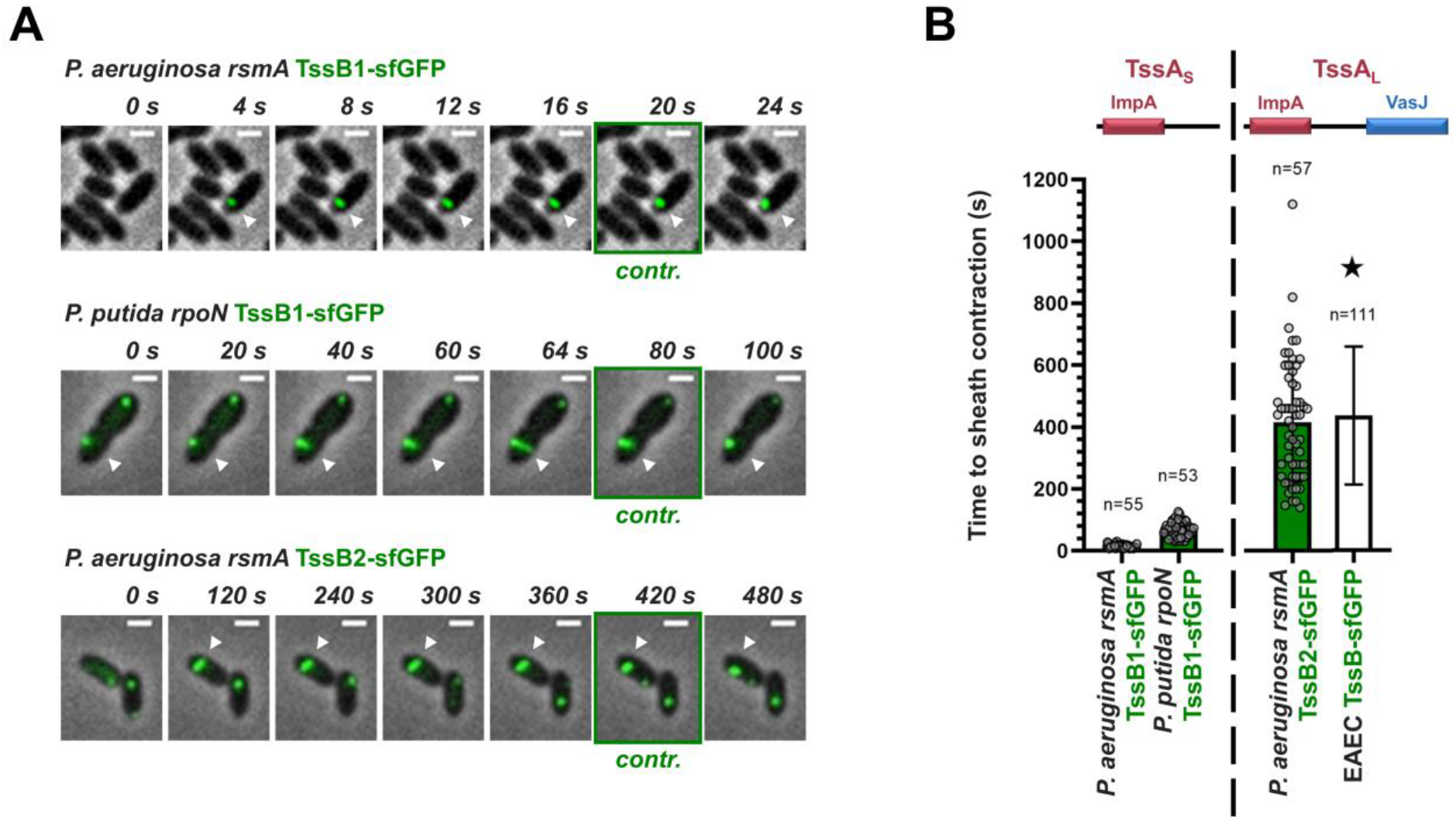
Distinct anchoring modes correlate with different T6SS behaviors. **(A)** T6SSs with short TssA proteins and TagB or TagJ anchors contract immediately after sheath extension; examples presented here are the *P. aeruginosa* H1-T6SS (TagJ-mediated sheath stabilization) and the *P. putida* K1-T6SS (TagB-mediated sheath stabilization) (top and middle, respectively). T6SSs with long TssA proteins tend to reside for prolonged periods of time in the cell after sheath extension. This is usually supported by TagA anchors (29, 30, 33), nonetheless for the example presented here (*P. aeruginosa* H2-T6SS; bottom), it has been proposed that TssA2 could act as an anchor itself (29), as no TagA protein has been identified for the H2-T6SS (File S1). The presented panels are selected images from fluorescence microscopy time-lapse recordings of *P. putida rpoN* expressing TssB1-sfGFP from the native *tssB1* locus and *P. aeruginosa rsmA* expressing TssB1-sfGFP and TssB2-sfGFP from the native *tssB1* and *tssB2* loci, respectively. White arrowheads indicate the T6SS sheaths of interest and panels showing sheath contraction (“contr.”) are marked with a green outline. Images were recorded every 2 s for the H1-and K1-T6SS and every 20 s for the H2-T6SS. Scale bars represent 1 μm. **(B)** Quantification of (A) and integration of the data collected as part of this study with measurements from the literature. Time to sheath contraction was measured for the T6SSs shown in panel (A); measurements were performed manually on fluorescence microscopy time-lapses of the relevant strains using the frame directly prior to that in which sheath contraction was observed. Values for the T6SS of enteroaggregative *E. coli* (EAEC) (30) (marked by a star) refer to residence times only. n indicates the number of cells that are included in the analysis. Whilst comprehensive measurements for the time to contraction of the *V. cholerae* T6SS, containing a TssA_L_ and a TagA_V_ anchor (not included in the graph), have not been performed, residence times of up to 220 s have been reported (29). Data collection and image analysis protocols for all panels presenting the results of fluorescence microscopy experiments are described in detail in the Materials and Methods.

## DISUSSION

In this study we have discovered and characterized a new T6SS stabilization mode that is employed by apparatuses containing short TssA proteins. This mode of sheath stabilization relies on the TssA-dependent recruitment of a novel class of protein partners (TagB or TagJ) to the baseplate structure. These T6SS components do not necessarily share any sequence similarity, but have near identical molecular weights (Fig. 1B). Our findings provide much needed insight into the anchoring of TssA_S_-containing T6SSs and allow us to propose a framework for T6SS sheath stabilization (Fig. 5). TssA_L_-containing T6SSs usually rely on membrane-associated TagA components for anchoring. TagA proteins interact with TssA_L_, stably clamping the end of the sheath at the opposite side of the cell and maintaining the sheath in the extended conformation (Fig. 5, top). Absence of TagA allows the end of the sheath to slide and the structure to keep elongating, something that delays contraction even further and often results in broken or floating sheaths of unnatural length (29, 30, 33). In contrast, T6SSs with TssA_S_ proteins have no terminal sheath anchor and are instead stabilized by the recruitment and accumulation of TagB or TagJ structural components at the baseplate (Fig. 3AB). One can envisage that these support proteins form a collar-like structure that constricts the baseplate (Fig. 3AB), preventing the conformational change (dome-to star-shaped baseplate) that leads to contraction (35) until the sheath polymerizes to its full size (Fig. 2DEF). The sheath contracts as soon as it reaches the opposing cell membrane (Fig. 1D and 3B), dispersing the TagB or TagJ multimer (Fig. 5, bottom). Both modes of sheath stabilization are important for T6SS efficacy in interbacterial killing, as loss of either type of stabilizing protein leads to increased prey survival (30) (Fig. 3DE). For TagA-mediated anchoring, decreased prey killing is due to unchecked polymerization and breaking of the sheaths (30). By contrast, for TagB1-mediated stabilization, the production of overly short sheaths results in a reduction in killing (Fig. 2DE and 3DE). This may be due to the delivery of fewer Hcp-associated effectors (lower *P. putida* Hcp1 secretion; Fig. 3C), reduced mechanical energy of the sheath, or a combination of both.

**Fig. 5.**
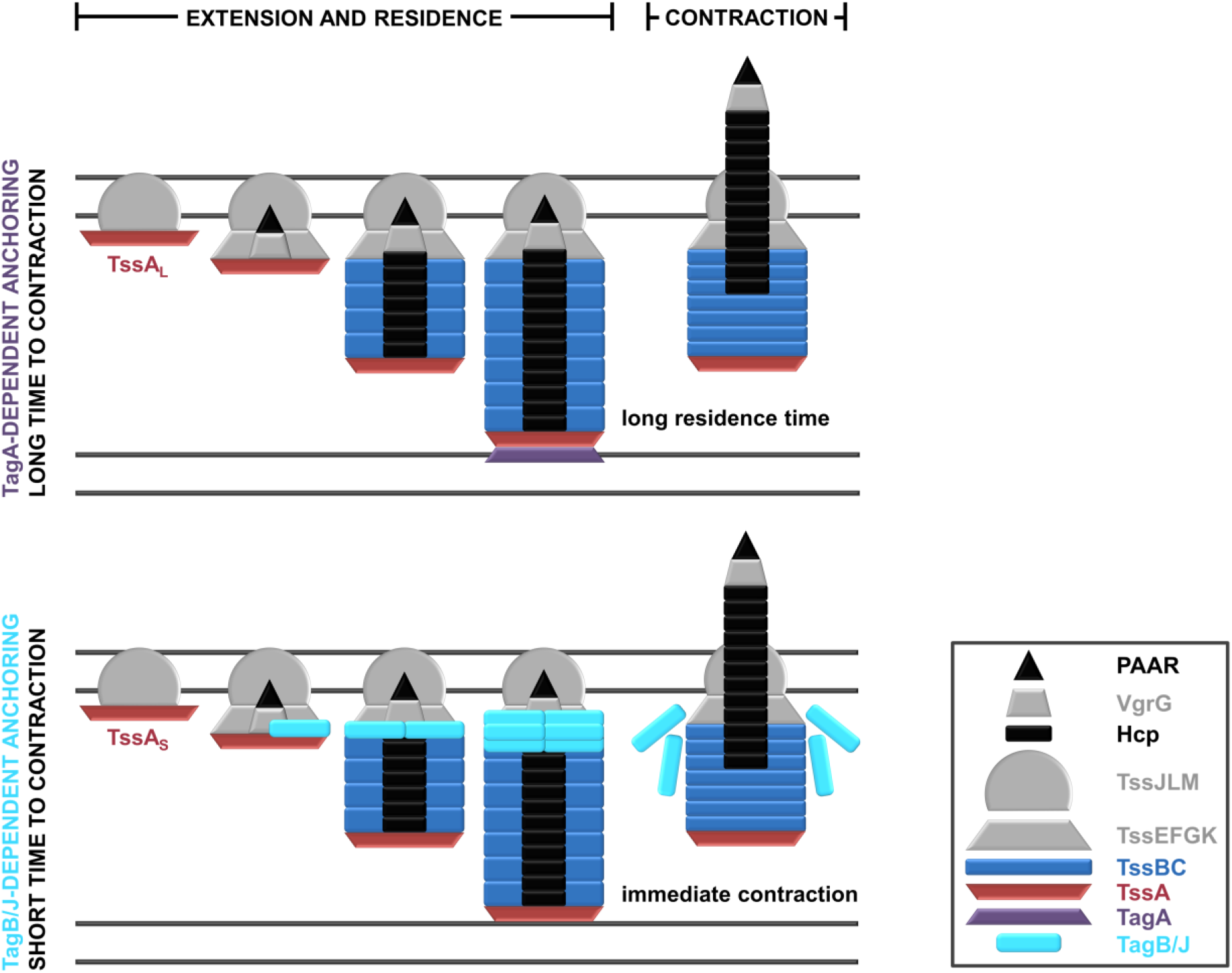
Schematic representation of the characterized modes of T6SS anchoring. **(Top)** T6SSs containing TssA_L_ proteins are usually anchored by TagA structural components. The TagA anchor is recruited by TssA_L_ as the latter approaches the opposing cell membrane and clamps the end of the sheath, stabilizing it for prolonged periods of time. **(Bottom)** T6SSs containing TssA_S_ proteins are stabilized by protein partners like TagB or TagJ, which are recruited to the baseplate by TssA. As the sheath extends with TssA at its end, TagB/TagJ continues to accumulate on the baseplate, maintaining this T6SS structure in a stable conformation and allowing the sheath to reach its full size. In this anchoring mode, T6SS firing occurs immediately after the sheath reaches the opposing cell membrane. The inset provides a key for the colors and shapes used to depict T6SS protein components.

We have shown that the pairing of different forms of TssA (TssA_L_ or TssA_S_) with diverse anchoring partners (TagA or TagB/J) generates distinct T6SS stabilization modes which correlate with longer or shorter times to sheath contraction (Fig. 4). Varied timescales of sheath assembly and firing naturally result in very different numbers of assembled T6SS apparatuses per cell. In cases where the sheath is stabilized via a TssA_S_-TagB/J mechanism, the vast majority of cells contain a single sheath at any given time, whereas for systems anchored by TssA_L_-TagA, cells often contain more than one sheath (29, 32) and can harbor up to six fully assembled structures (Fig. S7). Such diverse numbers of T6SSs per cell can generate varied aggression strategies. For T6SSs with short residence times, cells can engage in constant firing (*P. putida* K1-T6SS (31)) or quickly retaliate against incoming attacks (*P. aeruginosa* H1-T6SS (34)), whilst for T6SSs with long residence times and multiple apparatuses per cell, attackers can deploy a larger-than-normal effector payload upon a specific firing cue.

The description of T6SS sheath anchoring by TagB or TagJ proteins raises the question of how does sheath stabilization occur in T6SSs containing a TssA_S_ protein, but no identified anchor, for example T6SSs belonging to phylogenetic group 4A (Fig. 1B) like the *P. aeruginosa* H3-T6SS. To begin to probe this, we generated a *P. aeruginosa rsmA* strain expressing TssB3-sfGFP from the native *tssB3* locus and imaged the H3-T6SS sheaths. We found sheath assembly in this system to be markedly different to many of the T6SSs visualized previously. TssB3-sfGFP foci and assembled sheaths were always located at the cell pole (Fig. S8A), something that is not observed for the *P. aeruginosa* H1- and H2-T6SSs (Fig. S8B). H3-T6SS sheaths contract (Fig. S8C), nonetheless, instead of extending across the short axis of the cell like other sheaths imaged in this study, they extend from the cell pole, either across the long axis of the cell, leading to apparatuses with a “floating” end (Fig. S8A, red arrowhead), or diagonally, eventually leaning against the opposing cell wall (Fig. S8A, yellow arrowhead). This distinct behavior seems to be in agreement with the lack of an obvious sheath anchor (Fig. 1B) and alludes to a third mode of sheath stabilization that correlates with polar localization of the T6SS apparatus. It is likely that contraction of these sheaths will propel the Hcp3 tube to the exterior of the cell with very little mechanical force. Nonetheless, as no toxins have been associated with this system to date (the H3-T6SS has only been shown to secrete an effector involved in iron acquisition (6)), it is likely that very little force is needed, if the function of the H3-T6SS is to allow the secretion of a common-goods effector to the extracellular milieu, rather than puncturing prey cells to deliver toxins. In addition, the polar localization of the system possibly allows it to be poised and ready to fire when iron becomes limited, without obstructing the assembly of the other two T6SSs of this bacterium (36) which are usually not localized at the pole (Fig. S7A).

Overall, the distinct aggression behaviors that may emerge from the different modes of T6SS stabilization described here could benefit bacteria in different competition scenarios and ecological niches. This suggests that the functional diversity of T6SS apparatuses does not only stem from the broad range of effectors they can deliver (37, 38), but can also be generated by differences in the structural components of the system which underpin the modes and dynamics of T6SS assembly and firing.

## MATERIALS AND METHODS

A detailed description of the materials and methods used to generate the data supporting the findings in this study can be found in the Supplementary Information.

## ACKNOWLEDGEMENTS

We thank T. Ellis and C. Ramos for the kind gifts of the mScarlet fluorophore and *Pseudomonas savastanoi* pv. savastanoi strain NCPPB 3335, respectively, and we are grateful to all members of the Filloux laboratory for helpful discussions. This study was supported by the MRC grants MR/N023250/1 and MRK/K001930 (to A.F.), the BBSRC grant BB/N02539/1 (to A.F.), the MRC Career Development Award MR/M009505/1 (to D.A.I.M.), an institutional MRC studentship (to S.F), as well as the European Commission Marie Curie Fellowship H2020-MSCA-IF-2014-654135, the InterTalentum Fellowship GA713366 (co-funded by the European Commission and the Universidad Autónoma de Madrid), and the “Research Challenges” 2018 R+D+i Project RTI2018-096936-J-I00 (funded by the Spanish Ministry of Science, Innovation and Universities) (to P.B). The Facility for Imaging by Light Microscopy (FILM) at Imperial College London is part-supported by the Wellcome Trust grant 104931/Z/14/Z and the BBSRC grant BB/L015129/1.

## AUTHOR CONTRIBUTIONS

P.B., R.C.D.F., D.A.I.M. and A.F. designed the research. P.B. carried out *in silico* analyses and performed molecular biology and microbiology experiments. R.C.D.F. performed molecular biology, biochemistry and fluorescence microscopy experiments and analyzed fluorescence microscopy and mass spectrometry data. S.F constructed plasmids and *P. aeruginosa* strains. R.C.Y.L. performed competition assays with plant pathogens. L.S. performed bacterial-two-hybrid assays and constructed the associated plasmids. D.A.I.M performed protein purification and biochemistry experiments. P.B., R.C.D.F, D.A.I.M and A.F. wrote the manuscript with input from all authors. D.A.I.M. and A.F. directed the project.

