## Supplementary Information for "Novel structural components generate distinct type VI secretion system anchoring modes"

**This PDF file includes:**

- Supplementary Materials and Methods
- Figures S1 to S9
- Tables S1 to S3
- Legends for Videos S1 and S2
- Legends for Files S1 and S2
- Supplementary references

### SUPPLEMENTARY MATERIALS AND METHODS

***Reagents and bacterial growth conditions.*** Unless otherwise stated, chemicals, antibiotics and reagents were acquired from Sigma Aldrich and growth media were purchased from Merck or Oxoid. Lysogeny broth (LB) (10 g/L NaCl) and agar (1.5% w/v) were used for routine growth of all organisms with shaking at 200 RPM as appropriate; *E. coli* and *P. aeruginosa* were grown at 37 ºC, *P. putida* at 30 ºC and plant pathogens (*X. campestris*, *A. tumefaciens*, *P. syringae* and *P. savastanoi*) at 28 ºC. For competition assays with plant pathogens LB (5 g/L NaCl) was used, whereas microscopy experiments and secretion assays were performed using tryptone soya broth (TSB) (Oxoid). Growth media were supplemented with the following, as required: 1 mM Isopropyl β-D-1-thiogalactopyranoside (IPTG), 30 μg/mL tetracycline, 100 μg/mL ampicillin, 50 μg/mL kanamycin, 20 μg/mL gentamicin, 30 μg/mL chloramphenicol, 100-200 μg/mL streptomycin for *P. putida* and 2000 μg/mL streptomycin for *P. aeruginosa*, 25 μg/mL piperacillin, and 20 μg/mL rifampicin.

***Construction of plasmids and bacterial strains.*** Bacterial strains, plasmids and oligonucleotides used in this study are listed in Tables S1, S2 and S3, respectively. DNA manipulations were conducted using standard methods. KOD Hot Start DNA polymerase (Merck) was used for all PCR reactions according to the manufacturer's instructions, oligonucleotides were synthesized by Sigma Aldrich and restriction enzymes were purchased from Roche and New England Biolabs. All constructs were DNA sequenced and confirmed to be correct before use. Recombinant plasmids were transferred to *E. coli* strains by transformation and to *Pseudomonas* strains by electroporation (1) or conjugation (2), as appropriate.

Genes encoding T6SS structural components (*P. putida* Hcp1, TssA1, TagB1, TssE1, TssF1, TssG1, TssK1 and VgrG1 as well as *P. aeruginosa* TssA1 and TagJ1) were amplified from genomic DNA extracted from *P. putida* KT2440R and *P. aeruginosa* PAO1. *P. putida hcp1* was cloned into the IPTG-inducible plasmid pET28a using primers P1 and P2; when expressed, *P. putida* Hcp1 has an N-terminal $His_6$ tag. *P. putida tssA1* and *tagB1* and *P. aeruginosa tssA1* and *tagJ1* were cloned into the IPTG-inducible plasmid pETDuet-1, in MCS-1 (*tssA1*) and MCS-2 (*tagB1* and *tagJ1*) using primers P3-P18. When expressed, *P. putida* and *P. aeruginosa* TssA1 proteins have an N-terminal $His_6$ tag (genes cloned with primer pairs P5/P6 and P9/P10, respectively) or an N-terminal V5 tag (GKPIPNPLLGLDST) (primer pairs P11/P12 and P15/P16, respectively were used for exchange of the affinity tag). When expressed, TagB1 and TagJ1 have a C-terminal StrepII tag (genes cloned with primer pairs P3/P4 and P7/P8, respectively) or V5 tag (primer pairs P13/P14 and P17/P18, respectively were used for exchange of the affinity tag). *P. putida tssE1*, *tssF1*, t*ssG1*, *tssK1*, *hcp1*, *vgrG1* and *tssA1* were cloned into the bacterial-two-hybrid vector pKNT25 using primers P68-P80 in order to generate chimeric genes encoding N-terminal fusions of the respective proteins with the T25 fragment of *Bordetella pertussis* adenylate cyclase. *P. putida tagB1* was cloned into vectors pUT18C (primer pair P64/65) and pUT18 (primer pair P66/67) to produce chimeric genes encoding C- and N-terminal fusions of TagB1 with the T18 fragment of *B. pertussis* adenylate cyclase.

*P. putida* gene mutants (*tagB1* and *hcp1*) were constructed by allelic exchange, as previously described (3). Briefly, 500-bp DNA fragments upstream and downstream the gene to be deleted were amplified using *P. putida* KT2440 genomic DNA. A fragment containing both regions was obtained by overlapping PCR, cloned into pCR-BluntII-TOPO (Invitrogen), sequenced and subcloned into the *Xba*I/*Bam*HI sites of pKNG101 (primers P19-P22 (for *tagB1*) and P92-P95 (for *hcp1*)). The suicide vector pKNG101 (4) does not replicate in *Pseudomonas*; it was maintained in *E. coli* CC118λpir and mobilized into *Pseudomonas* by triparental conjugation.

A similar approach was used to replace wild-type *P. putida tssA1*, *tssB1*and *tagB1* as well as *P. aeruginosa tssB1*, *tssB2*, *tssB3* and *tagJ1* with genes encoding the protein of interest C- or N-terminally fused to a sfGFP or mScarlet-I fluorophore (all fusions carry the fluorophore at the C-terminus of the protein with the exception of the *P. putida* TssA1 fusion where the sfGFP is at the N-terminus of the protein); primers P23-P43, P46-P63 and P82-P87 were used for engineering these substitutions. The same strategy was used to replace the wild-type *P. putida tagB1* gene with a version encoding TagB1 C-terminally fused to two consecutive StrepII tags (primers P88-P91). The *tssB1*-mScarlet-I gene (amplified with primer pair P44/P45), encoding a C-terminal fusion of m-Scarlet-I to *P. putida* TssB1, was introduced on the chromosome using the miniCTX vector (5). All insertions and gene replacements were confirmed by PCR and DNA sequencing.

***Protein identification by mass spectrometry.*** Bacterial cultures of *P. putida rpoN* and of a *P. putida rpoN* strain where *tagB1* was replaced with a version of the gene encoding TagB1 C-terminally fused to two consecutive StrepII tags, were grown in TSB supplemented with the appropriate antibiotics for at least 8 hours at 30 ºC with shaking at 200 RPM. Bacterial suspensions were then sub-cultured at an $OD_{600}$ of 0.1 into 50 ml TSB and incubated for an additional 8 hours under the same growth conditions. Cells were harvested and cell pellets were resuspended in 50 mM Tris-HCl (pH 7.5), 150 mM NaCl and lysed by sonication after the addition of protease inhibitors (Roche). Cell debris was eliminated by centrifugation (48,000 *x* g, 30 mins, 4 °C) and both suspensions were subjected to protein purification using Strep-Tactin Sepharose (Iba Lifesciences), according to the manufacturer’s specifications. The resulting protein mixtures were methanol precipitated and freeze dried; three samples were prepared for each purification condition.

Freeze-dried protein mixtures were resuspended to a final concentration of 1 μg/mL in 25 μL of 20 mM ammonium bicarbonate before overnight digestion with trypsin at 30 °C (ratio 20:1 protein mixture:trypsin). Direct analysis of peptide mixtures was carried out under trap and elute conditions using an Acclaim Pepmap 100 Nano-Trap and column (Thermo Fischer Scientific) on an Eksigent nanoLC Ultra 2D HPLC coupled to a Triple-TOF 5600+ mass spectrometer via a Nanospray III source (all AB Sciex). The peptides were loaded in 98% v/v water, 2% v/v acetonitrile, 0.05% v/v trifluoroacetic acid and washed on the trap for 20 minutes, before being injected onto the column and eluted over 100 minutes with a rising gradient of acetonitrile composed of solutions A and B (solution A = 98% v/v water, 2% v/v acetonitrile, 0.1% v/v formic acid; solution B = 98% v/v acetonitrile, 2% v/v water, 0.1% v/v formic acid) as follows: 0 minutes, 98%A:2%B; 60 minutes, 80%A:20%B; 75 minutes, 60%A:40%B; 80 minutes, 2%A:98%B, 87.5 minutes, 98%A:2%B, 100 minutes, 98%A:2%B. Mass spectra were acquired between 400-1250 *m/z* and the top 20 peptides with a charge between +2 and +5 were selected to proceed to MS/MS (95-1800 *m/z*). All mass spectrometry was performed at the BBSRC Mass Spectrometry and Proteomics Facility at the University of St. Andrews, UK.

<u>*MaxQuant analysis*</u>. Data were processed using MaxQuant version 1.5.8.3 (6). Peptides were identified from MS/MS spectra searched against the Uniprot *P. putida* ATCC47054/DSM6125/NCIMB11950/KT2440 reference proteome (proteome ID: UP000000556) (accessed November 2019) using the Andromeda search engine (7).

Carbamidomethylation, methionine oxidation and N-terminal acetylation were specified as variable modifications. *In silico* digest of the reference proteome was performed using the Trypsin/P setting with up to two missed cleavages allowed. The false discovery rate (FDR) was set at 0.01 for peptides, proteins and sites. The "re-quantify" function was enabled. The sequence decoy mode used was "revert". Protein quantification was performed using the MaxLFQ algorithm within MaxQuant (8). Unique and razor peptides were used for quantification. All other parameters were used as pre-set in MaxQuant.

*Perseus analysis*. Data were analyzed using Perseus version 1.5.8.5 (9). Proteins present in the "reverse", "only identified by site" and "potential contaminant" databases were removed, and proteins identified by one or more unique peptides retained for further analysis. LFQ intensities were logarithmized ($\log_2$) and replicates grouped together before the data was filtered to retain only proteins identified in two or more replicates from at least one sample. Missing $\log_2$ LFQ intensities were inferred using a downshifted normal distribution (1.8 downshift, 0.3 width) and proteins that were significantly enriched by TagB1-(StrepII)$_2$ identified using a two-sample *t*-test (permutation based false discovery rate (FDR) = 250, FDR = 0.05, $S_0$ = 0.2).

***Far-western dot blots.*** *P. putida* TagB1 and *P. aeruginosa* TagJ1 self-interaction, as well as interaction of these proteins with their cognate TssA1 proteins were assessed using far-western dot blots as follows: *E. coli* BL21(DE3) cells expressing TagB1-StrepII, TagJ1-StrepII or a leaderless version of the *E. coli* cytochrome *c* maturation protein CcmG fused to a C-terminal StrepII tag (binding control protein) were grown in LB (10 g/L NaCl) supplemented with the appropriate antibiotics at 37 °C with shaking at 200 RPM to an $OD_{600}$ of 0.8. Expression was subsequently induced using IPTG for 16 hours at 30 °C. Cells were harvested and cell pellets were resuspended in 50 mM Tris-HCl (pH 7.5), 150 mM NaCl and lysed by sonication after the addition of protease inhibitors (Roche). Cell debris were eliminated by centrifugation (48,000 *x* g, 30 mins, 4 °C) and proteins were purified using Strep-Tactin Sepharose (Iba Lifesciences) according to the manufacturer's specifications. 3.5 ng of pure TagB1-StrepII, TagJ1-StrepII and CcmG-StrepII were spotted onto Amersham Protran nitrocellulose membranes (0.45 µm pore size, GE Life Sciences) and spots were dried at room temperature. Membranes were blocked overnight at 4 °C in 3% w/v Bovine Serum Albumin (BSA)/TBS-T (0.1% v/v Tween 20). The following day, 150 $OD_{600}$ units of *E. coli* BL21 (DE3) containing the empty vector (pETDuet-1) or over-expressing TagB1-V5, TagJ1-V5, *P. putida* V5-TssA1 or *P. aeruginosa* V5-TssA1 were resuspended in 10 mL of binding buffer (20 mM Tris-HCl (pH 8.0), 10% v/v glycerol, 100 mM NaCl, 3% w/v BSA) and lysed by sonication. Crude lysates, or 10 mL of binding buffer alone, were applied directly to the blocked nitrocellulose membranes before overnight incubation at 4 °C. Membranes were washed three times for 10 minutes with TBS-T and probed with mouse anti-V5 (Invitrogen) (dilution 1:5,000 in 3 w/v % BSA/TBS-T) or Strep-Tactin-HRP conjugate (Iba Lifesciences) (dilution 1:3,000 in 3 w/v % BSA/TBS-T) at room temperature for 4 hours. Goat anti-mouse IgG-HRP conjugate (Sigma Aldrich) (dilution 1:6,000 in 3% w/v BSA/TBS-T) was applied for 1 hour at room temperature, as appropriate. Membranes were washed three times for 10 minutes with TBS-T prior to development with Immobilon Classico Western HRP substrate (Merck Millipore) using a Gel DOC XR+ Imager (Bio-Rad).

***Interbacterial competition assays.*** *In vitro* competition assays were performed on LB (10 g/L NaCl) (*E. coli* prey) or LB (5 g/L NaCl) (plant pathogens as prey) agar (1.5% w/v) plates, as previously described (10). Briefly, overnight bacterial cultures were washed and adjusted to an $OD_{600}$ of 10 in sterile PBS, and mixed at a 1:1 ratio (*P. putida*:prey). Mixtures were grown on LB agar plates at 30 °C for 5 hours (*E. coli* prey) or 24 hours (plant pathogens as prey) and then collected using an inoculating loop and resuspended in sterile PBS. The outcome of the competition was quantified by counting colony forming units (CFUs) using antibiotic selection of the input (time = 0 hours) and output (time = 5 hours or time = 24 hours). All prey strains harboured the plasmid pRL662, which confers resistance to gentamicin and was used for antibiotic selection, whereas *P. putida* KT2440R is naturally resistant to rifampicin. *In planta* competition assays were carried out by infiltration of bacteria into *Nicotiana benthamiana* leaves, as described before (11). Briefly, overnight cultures of *P. putida* and *X. campestris* were adjusted to $OD_{600}$ of 0.1 in PBS and mixed at a 1:1 ratio. Approximately 100 µl of each suspension were infiltrated on the reverse of an approximately one-month-old leaf and the infiltration area was marked. After 24 hours of incubation in a plant chamber (23°C, 16 hours light exposure), a circular section of 6 mm diameter from the infiltration area of the leaf was isolated using a cork-borer set (Sigma-Aldrich), homogenized in PBS using a motorized tissue grinder (FisherBrand), serially diluted and platted on LB agar supplemented with the appropriate antibiotics. CFUs were determined using antibiotic selection as described above. For all competition assays, at least three biologically independent experiments were performed.

***P. putida Hcp1 antibody production:*** An overnight TSB culture of *E. coli* BL21(DE3) cells harboring a pET28a plasmid encoding *P. putida* Hcp1 with an N-terminal $His_6$ tag was used to inoculate 1L of TSB supplemented with kanamycin at an $OD_{600}$ of 0.1. The culture was incubated at 37 ºC with shaking at 200 RPM and protein expression was induced with 0.5 mM IPTG at an $OD_{600}$ of 0.5. Cultures were grown overnight at 18 ºC with shaking at 200 RPM. Cells were harvested and the cell pellet resuspended in 50 mM Tris-HCl (pH 7.5), 250 mM NaCl, 20 mM imidazole (buffer A) supplemented with protease inhibitors (Roche). The cells were lysed by sonication and cell debris were eliminated by centrifugation (48,000 *x* g, 30 mins, 4 °C). $His_6$-Hcp1 was purified by immobilized metal affinity chromatography using nickel-Sepharose resin (GE Healthcare) equilibrated in buffer A and eluted off the resin with buffer A containing 500 mM instead of 20 mM imidazole. The protein was subsequently subjected to gel filtration chromatography in 50 mM Tris-HCl (pH 7.5), 250mM NaCl using a prepacked Superdex 75 10/300 GL size-exclusion column (GE Healthcare). Polyclonal antibody production was carried out by Eurogentec through their speedy 28-Day program (2 rabbits; 4 protein injections and 3 bleedings per rabbit).

***Secretion assays.*** *P. putida* strains were grown in TSB for 8 hours at 30° C with shaking at 200 RPM and the extracellular fraction was obtained and analyzed as previously described (12). Briefly, the cell suspensions were spun three times at 10,000 *x g* for 20 minutes. Bacterial pellets were normalized and added directly to 1x Laemmli buffer, whilst the culture supernatants were collected and precipitated with trichloroacetic (TCA) acid overnight. Precipitants were then washed with acetone and resuspended in 1x Laemmli buffer. All samples were boiled for 15 minutes. SDS-PAGE analysis was carried out using 8% or 15% BisTris NuPAGE gels (ThermoFisher Scientific), MES/SDS running buffer prepared according to the manufacturer’s instructions and pre-stained protein markers (Kaleidoscope Prestained Standard, Bio-Rad). Proteins were transferred to Amersham Protran nitrocellulose membranes (0.45 µm pore size, GE Life Sciences) using a Trans-Blot Turbo transfer system (Bio-Rad) before blocking in 5% w/v skimmed milk/TBS-T and adding the primary and secondary antibodies. The following primary antibodies were used: rabbit anti-Hcp1 antibody (dilution 1:500 in 5 w/v % skimmed milk/TBS-T) and mouse *E. coli* anti-RNA polymerase beta (Neoclone) (dilution 1:5,000 in 5 w/v % skimmed milk/TBS-T). The following secondary antibodies were used: goat anti-rabbit IgG-HRP conjugate (Sigma Aldrich) (dilution 1:6,000 in 5% w/v skimmed milk/TBS-T) and goat anti-mouse IgG-HRP conjugate (Sigma Aldrich) (dilution 1:6,000 in 5% w/v skimmed milk/TBS-T). Membranes were washed three times for 5 minutes with TBS-T prior to development. HRP conjugates were visualized with the Luminata Forte Western HRP Substrate (Merck) using a Gel Doc XR+ Imager (Bio-Rad).

***Bacterial two-hybrid assay.*** Protein-protein interactions were analyzed using the bacterial-two-hybrid (BTH) approach (13). Briefly, the proteins to be tested were independently fused to the T18 or T25 catalytic domains of the *B. pertussis* adenylate cyclase using plasmids pUT18 (or pUT18C) and pKT25, respectively. The two plasmids expressing the fusion proteins were simultaneously introduced into the reporter strain BTH101 by transformation and the plates were incubated at 30 °C for 24 hours. Three independent colonies for each plasmid combination were inoculated into 1 ml of LB (10 g/L NaCl) supplemented with ampicillin, kanamycin and 0.5 mM IPTG. After overnight growth at 30 °C with shaking at 200 RPM, 10 µl of each culture were spotted onto LB (10 g/L NaCl) agar (1.5% w/v) plates supplemented with ampicillin, kanamycin, 0.5 mM IPTG and 40 µg/ml X-Gal and incubated at 30 °C for 16 hours. At least three biologically independent experiments were performed for each protein pair.

***Fluorescence microscopy.*** Fluorescence microscopy experiments were performed using sfGFP (14) and mScarlet-I (15) fusion proteins. The functionality of the T6SS in *P. putida rpoN* strains expressing fluorescent fusions of TssA1 and TagB1 from the native gene loci, was confirmed by performing secretion assays with the strains *P. putida rpoN* TssA1-sfGFP and *P. putida rpoN* TagB1-sfGFP, as described above (Fig. S9). Bacterial cultures were grown in TSB supplemented with the appropriated antibiotics for at least 8 hours at 30ºC (*P. putida*) or 37 ºC (*P. aeruginosa*) with shaking at 200 RPM. Bacterial suspensions were then sub-cultured at an $OD_{600}$ of 0.1 into 50 ml TSB and grown under the same conditions to an $OD_{600}$ of 5 before visualization; *P. aeruginosa* PAO1 *rsmA tssB3::sfGfp* was grown at 25 °C with shaking at 200 RPM for 12 hours before visualization.

To visualize cells, 5 cm diameter glass bottom Petri dishes with a 3-cm diameter uncoated n° 1.5 glass window (MatTek Corporation) were used. 6 µl of bacterial culture were placed onto a 2.8 cm diameter (0.5 cm thickness) circular slab of Vogel-Bonner salts agar (2% w/v) which was, in turn, placed face down onto the glass bottom window of the Petri dish, so that the cells were sandwiched between the glass and the solid medium. Samples were immediately transferred to the microscope and imaged using an Axio Observer Z1 (Zeiss) inverted widefield microscope equipped with a Plan-Apochromat 63x/1.4 NA Oil Ph3 M27 objective (Zeiss), a SpectraX LED light engine (Lumencore), an ORCA-Flash 4.0 digital CMOS camera (Hamamatsu) and an environmental control system. Images were acquired using 1x1 binning, corresponding to a pixel size of 0.103 x 0.103 µm. All microscopy was performed at the Facility for Imaging by Light Microscopy (FILM) at Imperial College London, UK.

For the visualization of *P. putida* strains expressing TagB1-sfGFP or TssB1-sfGFP, fluorescence images were acquired every 2 s for a total of 180 s using an exposure time of 195 ms. For *P. putida* simultaneously expressing TssB1-mScarlet-I and sfGFP-TssA1, fluorescence images were acquired every 4 s for a total of 180 s using an exposure time of 500 ms for sfGFP and 250 ms for mScarlet-I. For *P. putida* simultaneously expressing TssB1-mScarlet-I and TagB1-sfGFP, fluorescence images were acquired every 2 s for a total of 180 s using an exposure time of 200 ms for sfGFP and 350 ms for mScarlet-I. For the visualization of *P. aeruginosa* expressing TssB1-sfGFP, fluorescence images were acquired every 2 s for a total of 180 s using an exposure time of 80 ms. For *P. aeruginosa* expressing TssB2-sfGFP, fluorescence images were acquired every 20 s for a total of 1200 s using an exposure time of 80 ms; the microscope chamber was set to 30 °C and the Definite Focus autofocus system was used. For *P. aeruginosa* expressing TssB3-sfGFP, fluorescence images were acquired every 30 s for a total of 450 s using an exposure time of 80 ms. Finally, for *P. aeruginosa* expressing TagJ1-sfGFP, fluorescence images were acquired every 60 s for 300 s using an exposure time of 400 ms.

***Image analysis and presentation.*** Image processing and analysis was performed using FIJI (16) in conjunction with the MicrobeJ (17) plugin. Generation of images was performed using the Zeiss-associated ZEN software (ZEN Blue) and FIJI. Prior to analysis, x-y drift present in time-lapse series was corrected for each channel using the StackReg plug-in for FIJI and "rigid body" transformation (18). Photobleaching in time-lapse series was corrected using the "simple ratio" bleach correction method within FIJI. Semi-automatic particle detection, segmentation and generation of cell masks were performed on phase-contrast images using the "medial axis" cell contour method within MicrobeJ; all cell masks were manually checked before further analysis. Cell counts and cell width data were directly extracted from manually curated cell masks generated using this method. Detection of fluorescent *P. putida* TagB1 foci was performed on fluorescence images using the local maxima detection algorithm implemented in FIJI, in conjunction with the "foci" particle conversion method implemented in MicrobeJ. Enumeration, localization and sheath length analysis of TssB1-sfGFP sheaths was performed manually for all visualized strains. Sheath length measurements were performed on time-lapse series acquired using *P. putida* strains expressing TssB1-sfGFP; measurements were performed on the frame directly prior to that in which sheath contraction was observed, using a custom line/ROI measurement macro in FIJI. Time to sheath contraction data were extracted manually from time-lapse series covering full sheath polymerization and contraction events. For image presentation, frames of interest were extracted from time-lapse series, corrected as described above, and scale bars created using the Scale Bar function in FIJI. Videos were prepared from the same corrected images and encoded using QuickTime (H.264 codec) (Apple Inc.). All videos are presented at 7 frames/s as .mov files.

***Statistical analysis of experimental data.*** Unless otherwise stated, statistical analysis was performed in GraphPad PRISM v8.0.2 using an unpaired two-tailed T-test with Welch's correction, multiple unpaired T-tests corrected for multiple comparisons using the Holm-Sidak method, or a one-way ANOVA with Tukey's multiple comparisons test, as appropriate. Statistical significance was defined as $p < 0.05$. Detailed information for each figure is provided below:

<u>Fig. 2E</u>*:* unpaired T-test with Welch's correction; *P. putida rpoN* TssB1-sfGFP (n=98), *P. putida rpoN tagB1* TssB1-sfGFP (n=95); 174.8 degrees of freedom, t-value=16.71; p=5.065 x $10^{-38}$ (significance).
<u>Fig. 2F</u>*:* unpaired T-test with Welch's correction; *P. putida rpoN* TssB1-sfGFP (n=706), *P. putida rpoN tagB1* TssB1-sfGFP (n=693); 1389 degrees of freedom, t-value=0.2638; p=0.7920 (non-significance).
<u>Fig. 3D</u>*:* unpaired T-tests corrected for multiple testing using the Holm-Sidak method. Number of n=5; 24 degrees of freedom. For *E. coli* competitions: *P. putida* vs. *P. putida tssA1* adjusted p=0.000063, *P. putida* vs. *P. putida tagB1* adjusted p=0.026749, *P. putida tssA1* vs. *P. putida tagB1* adjusted p=0.007411 (all significant). For *X. campestris*

competitions: *P. putida* vs. *P. putida tssA1* adjusted p=0.005651, *P. putida* vs. *P. putida tagB1* adjusted p=0.005814, *P. putida tssA1* vs. *P. putida tagB1* adjusted p=0.007411 (all significant). For *A. tumefaciens* competitions: *P. putida* vs. *P. putida tssA1* adjusted p=0.004148, *P. putida* vs. *P. putida tagB1* adjusted p=0.026749, *P. putida tssA1* vs. *P. putida tagB1* adjusted p=0.028596 (all significant). For *P. syringae* competitions: *P. putida* vs. *P. putida tssA1* adjusted p=0.004148, *P. putida* vs. *P. putida tagB1* adjusted p=0.002842, *P. putida tssA1* vs. *P. putida tagB1* adjusted p=5.976 $x10^{-8}$ (all significant). For *P. savastanoi* competitions: *P. putida* vs. *P. putida tssA1* adjusted p=0.005651, *P. putida* vs. *P. putida tagB1* adjusted p=0.000896, *P. putida tssA1* vs. *P. putida tagB1* adjusted p=0.000628 (all significant).
*Fig. 3E:* one-way ANOVA with Tukey's multiple comparison test; n=4; 11 degrees of freedom; F value=0.2636; p=0.0002 (significance). *P. putida* vs. *P. putida tssA1* adjusted p=0.0002. *P. putida* vs. *P. putida tagB1* adjusted p=0.0090. *P. putida tssA1* vs. *P. putida tagB1* adjusted p=0.0354.

***In silico* analyses.** In order to categorize TssA-like protein and identify TssA-related accessory T6SS structural components, an *in silico* study of 100 T6SS clusters was performed. Proteins encoding the core components and accessory proteins of the selected T6SS clusters were downloaded from the SecReT6 website (19) and organized by phylogenetic groups (groups 1-5, with phylogenetic subgroups 4A and 4B being considered separately); 20 T6SS clusters per phylogenetic group were examined. The T6SS cluster that is best characterized from each phylogenetic group was set as the reference cluster for the analysis (distinct clusters from *P. aeruginosa, E. coli, P. putida* and *A. tumefaciens* were selected as references). The reference clusters were used as queries in a blastp search (20) against the remaining selected T6SS clusters from the same phylogenetic group. The percentage identity and coverage resulting from the search can be found in File S1. Through this analysis, five families of TssA-like proteins were identified, and their domains were further characterized using the "NCBI Conserved Domain" function and the SMART website (21).

***Data availability.*** All data generated during this study that support the findings are included in the manuscript or in the Supplementary Information. All materials are available from the corresponding authors upon request.

SUPPLEMENTARY FIGURES

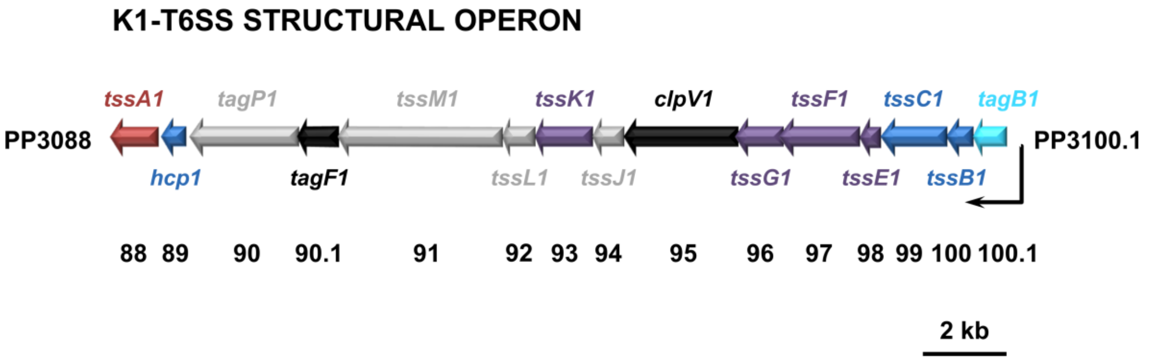


**Fig. S1. Schematic representation of the genomic organization of the structural genes of the K1-T6SS cluster from *P. putida* KT2440.** Genes encoding membrane complex proteins are shown in grey, whereas baseplate and tail components are depicted in purple and blue, respectively. *tssA1* is in red, *tagB1* in cyan and *tagF1* and *clpV1* are colored black. *vgrG1* is not shown, as it is located downstream in the effector module (22), which is omitted from this schematic.

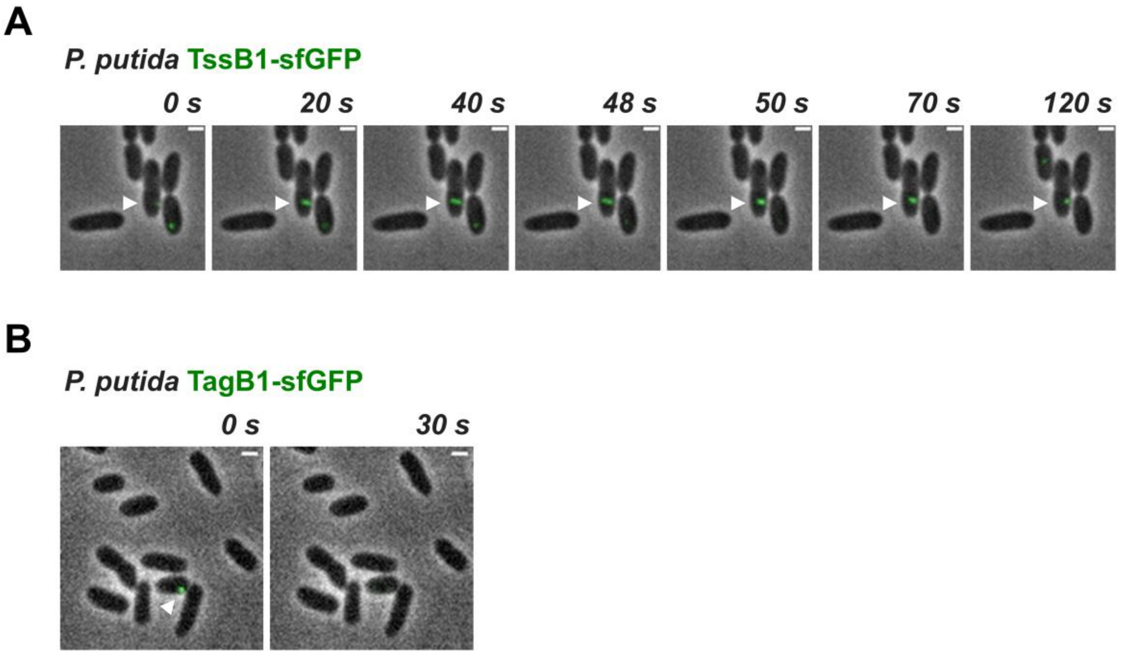


**Fig. S2. (AB) TssB1-sfGFP and TagB1-sfGFP show the same *in vivo* behavior in a wild-type *P. putida* strain as in *P. putida rpoN*. (A)** *In vivo* imaging of *P. putida* TssB1-sfGFP. Sheaths have identical dynamics in wild-type *P. putida* as observed in *P. putida rpoN*. The full cycle of T6SS assembly, including sheath initiation, extension, contraction and disassembly over approximately 120 seconds is shown for a representative sheath (white arrowhead). The presented panels are selected images from a fluorescence microscopy time-lapse recording of *P. putida* expressing TssB1-sfGFP from the native *tssB1* locus. Images were recorded every 2 s and scale bars represent 1 µm. **(B)** *In vivo* imaging of *P. putida* TagB1-sfGFP. TagB1 forms the same transient foci in wild-type *P. putida* as observed in *P. putida rpoN*; foci form over time and abruptly disappear (white arrowhead). The presented panels are selected images from a fluorescence microscopy time-lapse recording of *P. putida* expressing TagB1-sfGFP from the native *tagB1* locus. Scale bars represent 1 µm. Data collection and image analysis protocols are described in detail in the Materials and Methods.

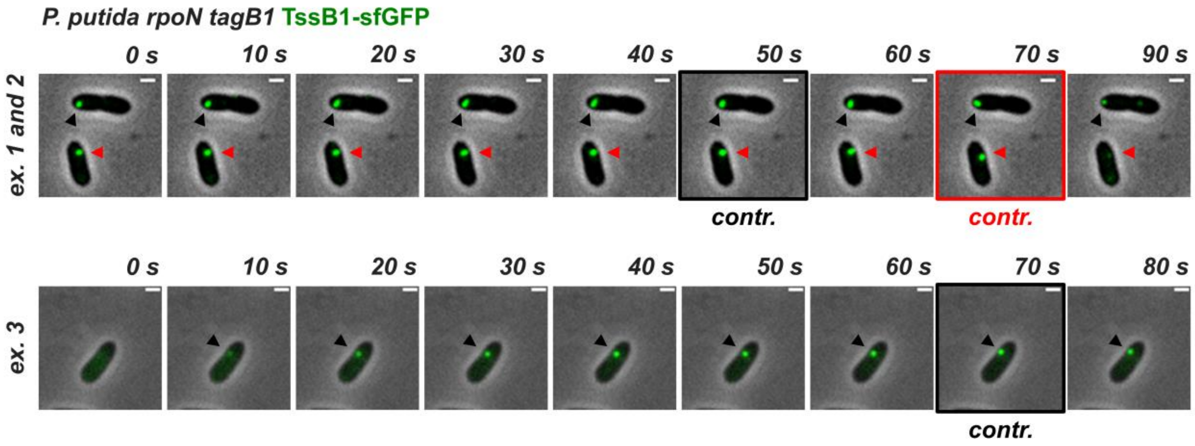


**Fig. S3. Absence of TagB1 prevents full extension of the K1-T6SS sheath.** *In vivo* imaging of *P. putida rpoN tagB1* TssB1-sfGFP. Three examples ("ex.") of T6SS sheaths that contract before fully extending to the opposite side of the cell are shown. Black or red arrowheads indicate the sheaths of interest and panels showing sheath contraction ("contr.") are marked with a back or red outline, respectively. The panels presented are selected images from two fluorescence microscopy time-lapse recordings of *P. putida rpoN* expressing TssB1-sfGFP from the native *tssB1* locus. Images were recorded every 2 s and scale bars represent 1 µm. An image from example 2 (top row; red arrowhead) was used to showcase the *P. putida rpoN tagB1* TssB1-sfGFP behavior in Fig. 2D. Data collection and image analysis protocols are described in detail in the Materials and Methods.

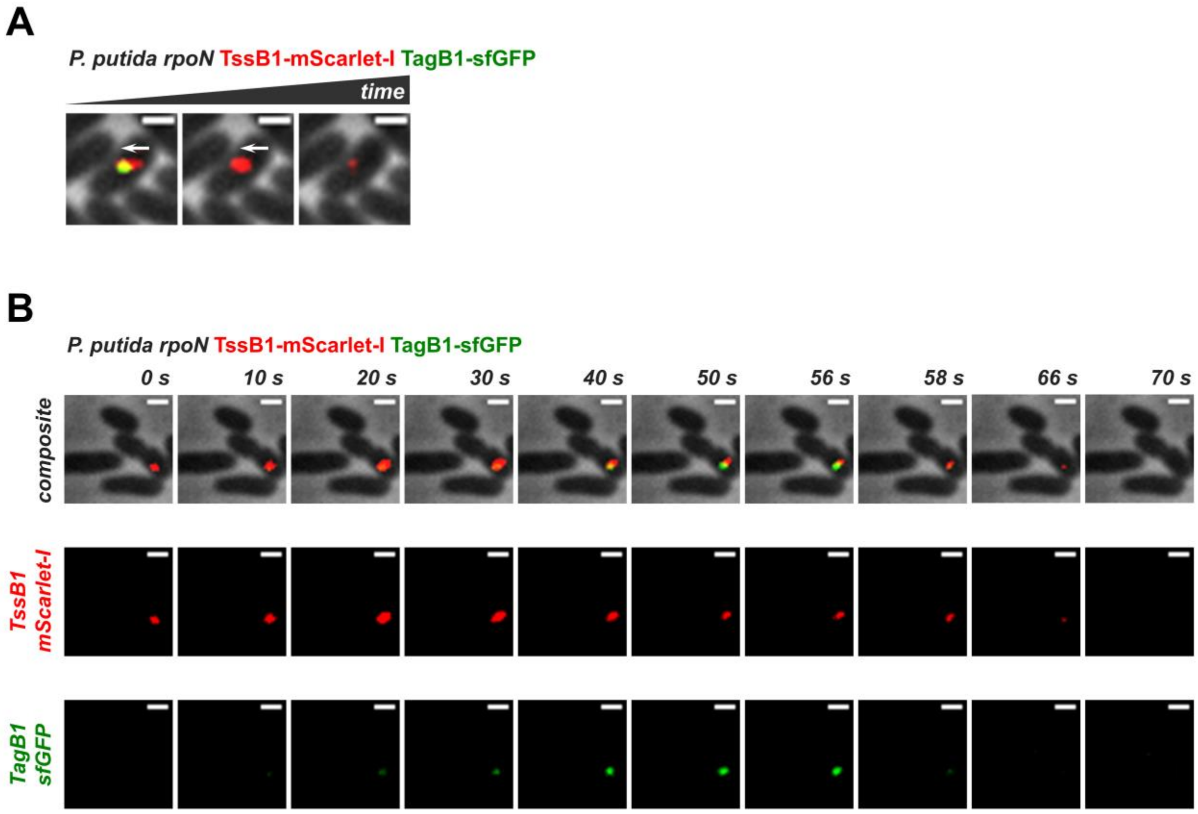


**Fig. S4.** ***P. putida*** **TagB1 localizes to the baseplate.** The panels presented are selected images from a fluorescence microscopy time-lapse recording of *P. putida rpoN* expressing TagB1-sfGFP from the native *tagB1* locus and the gene encoding TssB1-mScarlet-I integrated into the chromosome using the miniCTX plasmid (5) whilst the native *tssB1* copy is still present. Scale bars represent 1 µm. **(A)** TagB1-sfGFP foci are associated with the end of the extended sheath towards which contraction occurs. The white arrow indicates the direction of sheath contraction. **(B)** Images used to generate Fig. 3B. Phase-contrast composite images displaying both fluorescence channels are shown in the top row, whilst images from individual fluorescence channels are shown in the middle and bottom rows (mScarlet-I and sfGFP, respectively). Data collection and image analysis protocols for all panels presenting the results of fluorescence microscopy experiments are described in detail in the Materials and Methods.

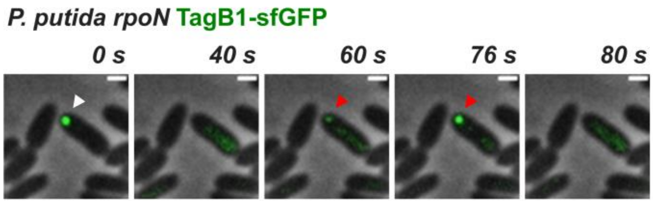


**Fig. S5.** ***P. putida*** **TagB1-sfGFP foci re-appear in the same location after a contraction event.** A TagB1-sfGFP focus (white arrowhead) disappears, presumably after sheath contraction, and a new focus appears (red arrowhead) in exactly the same location. The presented panels are selected images from a fluorescence microscopy time-lapse recording of *P. putida rpoN* expressing TagB1-sfGFP from the native *tagB1* locus. Images were recorded every 2 s and scale bars represent 1 µm. Data collection and image analysis protocols are described in detail in the Materials and Methods.

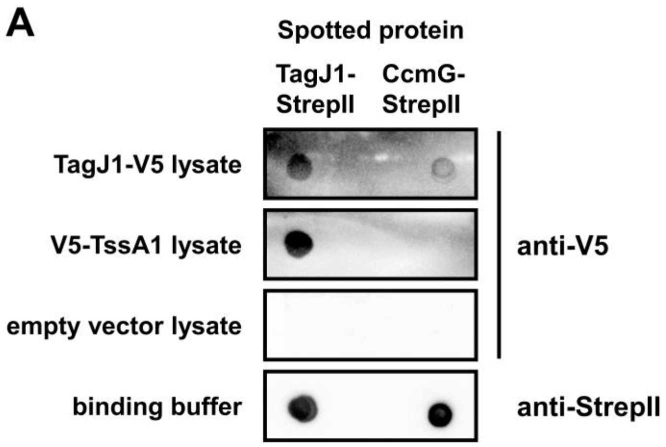


**Fig. S6. *P. aeruginosa* TagJ1 exhibits similar properties and *in vivo* behavior as *P. putida* TagB1. (A)** Far-western dot blot assays showing interaction of *P. aeruginosa* TagJ1 with itself and its cognate TssA1. Purified TagJ1-StrepII was spotted on a nitrocellulose membrane and was exposed to *E. coli* cell lysates expressing TagJ1-V5 or V5-TssA1. The TagJ1 interactions are specific as no interactions with the tested lysates are detected when the binding control protein CcmG-StrepII is probed or when TagJ1-StrepII is exposed to lysates from cells harboring the empty vector; similar amounts of TagJ1-StrepII and CcmG-StrepII were spotted on the membrane. Levels of StrepII-tagged proteins were assessed using a Strep-Tactin-HRP conjugate and levels of V5-tagged proteins were detected with an anti-V5 antibody and an HRP-conjugated secondary antibody. **(B)** *In vivo* imaging of *P. aeruginosa* TagJ1-sfGFP. TagJ1 forms transient foci which abruptly disappear (black arrowheads). Disappearance of foci is not due to photobleaching as new foci appear in directly adjacent cells (red arrowhead). The presented panels are selected images from a fluorescence microscopy time-lapse recording of *P. aeruginosa rsmA* expressing TagJ1-sfGFP from the native *tagJ1* locus. Images were recorded every 2 s and scale bars represent 1 µm. Data collection and image analysis protocols are described in detail in the Materials and Methods.

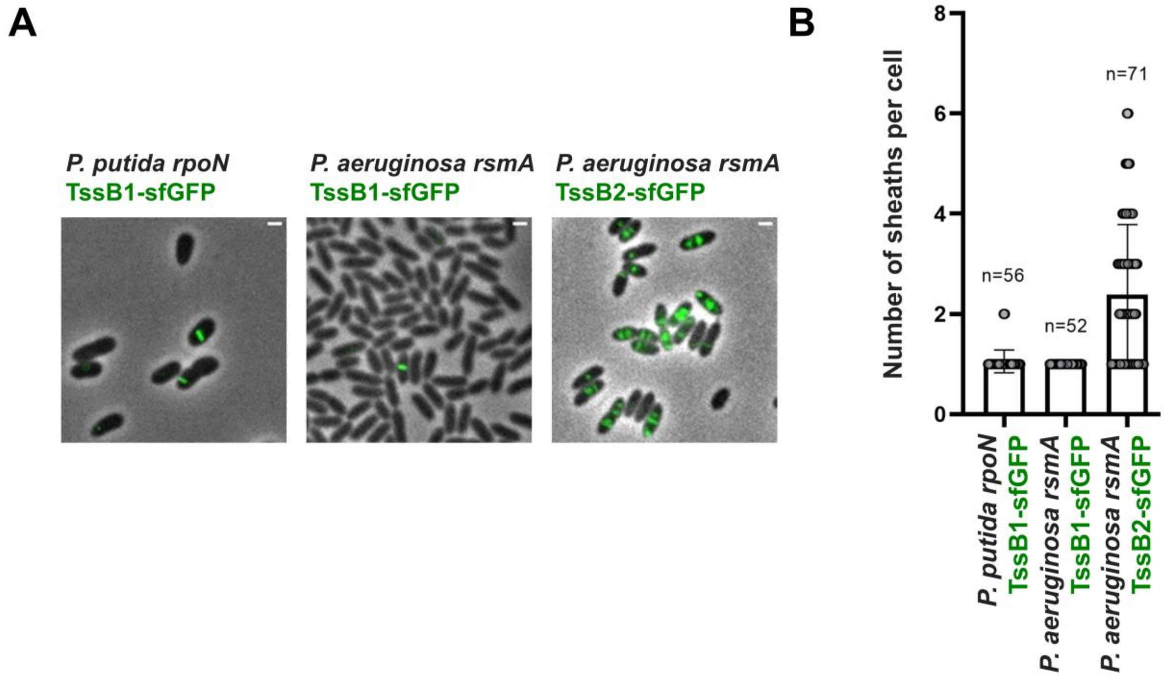


**Fig. S7. Distinct anchoring modes result in different numbers of T6SS apparatuses per cell. (A)** Representative fluorescence microscopy images of *P. putida rpoN* expressing TssB1-sfGFP from the native *tssB1* locus (left), and *P. aeruginosa rsmA* expressing TssB1-sfGFP (middle) and TssB2-sfGFP (right) from the native *tssB1* and *tssB2* loci, respectively. Sheath stabilization involving $TssA_S$ and TagB/J proteins leads to the presence of one T6SS sheath per cell, whereas $TssA_L$/TagA-mediated anchoring results in the co-existence of multiple sheaths per cell (23, 24). Scale bars represent 1 µm. **(B)** Quantification of (A). One apparatus is usually present at any given time for the *P. putida* K1-T6SS and *P. aeruginosa* H1-T6SS, but up to six sheaths can be simultaneously detected for *P. aeruginosa* H2-T6SS. n indicates the number of cells included in the analysis. Data collection and image analysis protocols for all panels presenting the results of fluorescence microscopy experiments are described in detail in the Materials and Methods.

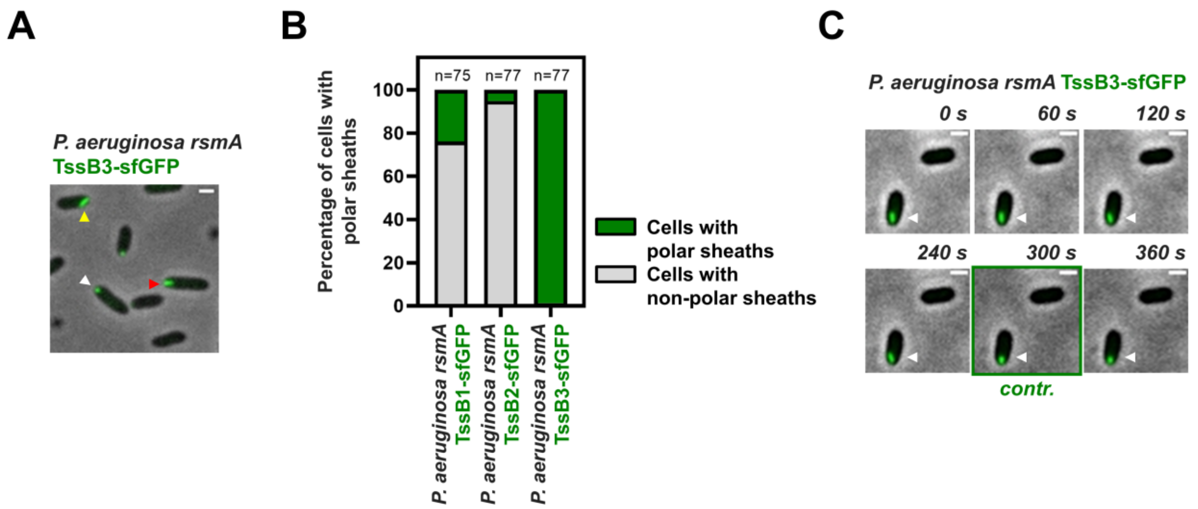


**Fig. S8. The *P. aeruginosa* H3-T6SS is not anchored in a classical manner. (A)** Representative fluorescence microscopy image of *P. aeruginosa rsmA* expressing TssB3-sfGFP from the native *tssB3* locus. TssB3-sfGFP foci (white arrowhead) and H3-T6SS sheaths (red and yellow arrowheads) are exclusively localized at the cell pole. Sheath structures grow from the pole either horizontally without reaching the opposite side of the cell (red arrowhead) or diagonally eventually leaning against the side of the cell (yellow arrowhead). Scale bars represent 1 µm. **(B)** Quantification of sheaths with polar localization in the *P. aeruginosa* H1-, H2- and H3-T6SS. Whilst the majority of the H1- and H2-T6SS sheaths are not localized at the cell pole, all H3-T6SS sheaths show polar localization. n indicates the number of cells included in the analysis. **(C)** *P. aeruginosa* H3-T6SS sheaths contract. A "floating" polar H3-T6SS sheath (white arrowhead) contracts over time; contraction ("contr.") is marked with a green panel outline. The presented panels are selected images from a fluorescence microscopy time-lapse recording of *P. aeruginosa rsmA* expressing TssB3-sfGFP from the native *tssB3* locus. Images were recorded every 30 s and scale bars represent 1 µm. Data collection and image analysis protocols for all panels presenting the results of fluorescence microscopy experiments are described in detail in the Materials and Methods.

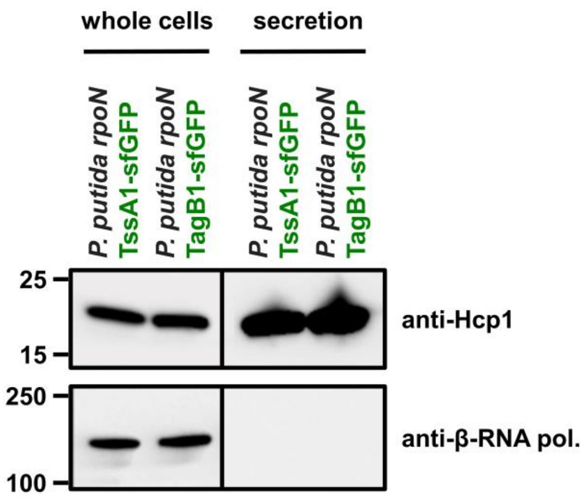


**Fig. S9. The functionality of the T6SS in *P. putida rpoN* strains expressing fluorescent fusions of TssA1 and TagB1 was confirmed by monitoring Hcp1 secretion.** Hcp1 expression and presence in the culture supernatant were assessed for *P. putida rpoN* strains expressing TssA1-sfGFP and TagB1-sfGFP from the native gene loci. Hcp1 protein levels were detected using an anti-Hcp1 primary antibody and an HRP-conjugated secondary antibody. Positions of molecular weight markers are shown on the left, the β-subunit of the *E. coli* RNA polymerase (β-RNA pol.) was used as a loading and bacterial lysis control and black lines indicate where the membrane was cut. A representative blot from three independent experiments is presented.

**SUPPLEMENTARY TABLES**

**Table S1.** Bacterial strains used in this study.

| Name | Description | Source |
|---|---|---|
| ***Escherichia coli*** | | |
| DH5α | F$^{-}$ *end*A1 *gln*V44 *thi*-1 *rec*A1 *rel*A1 *gyr*A96 *deo*R *nup*G *pur*B20 φ80d*lac*ZΔM15 Δ(*lac*ZYA-*arg*F)U169 *hsd*R17($r_K^- m_K^+$) λ$^{-}$ | (25) |
| CC118λpir | *ara*D Δ(*ara*, *leu*) Δ*lac*Z74 *pho*A20 *gal*K *thi*-1 *rsp*E *rpo*B *arg*E *rec*A1 λ*pir* | (26) |
| HB101 | *sup*E44 *hsd*S20 *rec*A13 *ara*-14 *pro*A2 *lac*Y1 *gal*K2 *rps*L20 *xyl*-5 *mtl*-1 | (27) |
| BL21(DE3) | F$^{-}$ *omp*T *gal dcm lon hsd*S$_B$($r_B^- m_B^-$) λ(DE3 [*lac*I *lac*UV5-T7p07 *ind*1 *sam*7 *nin*5]) [*mal*B$^{+}$]$_{K-12}$(λ$^{S}$) | (28) |
| BTH101 | F$^{-}$ *cya*-99 *ara*D139 *gal*E15 *gal*K16 *rps*L1(Str$^{R}$) *hsd*R2 *mcr*A1 *mcr*B1 | (13) |
| ***Pseudomonas putida*** | | |
| KT2440R | Rif$^{R}$ | (29) |
| KT2440R *tssA1* | Markerless mutant, Rif$^{R}$ | (22) |
| KT2440R *hcp1* | Markerless mutant, Rif$^{R}$ | This study |
| KT2440R *tagB1* | Markerless mutant, Rif$^{R}$ | This study |
| KT2440R *tagB1::sfgfp* | Rif$^{R}$ | This study |
| KT2440R *tssB1::sfgfp* | Rif$^{R}$ | This study |
| KT2440 *rpoN* | *rpoN*::*aphA*, Kan$^{R}$ | (30) |
| KT2440 *rpoN tssA1* | *rpoN*::*aphA*, Kan$^{R}$ | (22) |
| KT2440 *rpoN tssA1::sfgfp* | *rpoN*::*aphA*, Kan$^{R}$ | This study |
| KT2440 *rpoN tssA1 tssB1*::*sfgfp* | *rpoN*::*aphA*, Kan$^{R}$ | This study |
| KT2440 *rpoN tssA1 tagB1*::*sfgfp* | *rpoN*::*aphA*, Kan$^{R}$ | This study |
| KT2440 *rpoN tagB1* | *rpoN*::*aphA*, Kan$^{R}$ | This study |
| KT2440 *rpoN tagB1::sfgfp* | *rpoN*::*aphA*, Kan$^{R}$ | This study |
| KT2440 *rpoN tagB1::sfgfp tssB1-mScarlet-I* | *rpoN*::*aphA*, Kan$^{R}$, Tet$^{R}$ | This study |
| KT2440 *rpoN tagB1 tssB1*::*sfgfp* | *rpoN*::*aphA*, Kan$^{R}$ | This study |
| KT2440 *rpoN tssB1*::*sfgfp* | *rpoN*::*aphA*, Kan$^{R}$ | This study |
| KT2440 *rpoN tssB1*::*mScarlet-I* | *rpoN*::*aphA*, Kan$^{R}$ | This study |
| KT2440 *rpoN tssB1*::*mScarlet-I tssA1::sfgfp* | *rpoN*::*aphA*, Kan$^{R}$ | This study |
| KT2440 *rpoN tagB1*::*(StrepII)*$_2$ | *rpoN*::*aphA*, Kan$^{R}$ | This study |
| ***Pseudomonas aeruginosa*** | | |
| PAO1 *rsmA* | Markerless mutant | (31) |
| PAO1 *rsmA tssB1*::*sfgfp* | Markerless mutant | This study |
| PAO1 *rsmA tagJ1*::*sfgfp* | Markerless mutant | This study |
| PAO1 *rsmA tssB2*::*sfgfp* | Markerless mutant | This study |
| PAO1 *rsmA tssB3*::*sfgfp* | Markerless mutant | This study |
| **Plant pathogens** | | |
| *Xanthomonas campestris* pv. campestris IVIA 2734-1 | - | (22) |
| *Agrobacterium tumefaciens* C58 | - | (32) |
| *Pseudomonas syringae* pv tomato DC3000 | - | (33) |
| *Pseudomonas savastanoi* pv savastanoi strain NCPPB 3335 | - | (34) |

**Table S2.** Plasmids used in this study. Superscripts "PP" and "PA" indicate *P. putida* and *P. aeruginosa* components (genes or proteins), respectively; these abbreviations are solely used in this table for brevity.

| Name | Description | Source |
|---|---|---|
| pCR-BluntII-TOPO | Cloning vector, ColE1 *ori*, $Kan^R$ | Invitrogen |
| pETDuet-1 | Dual expression vector, ColE1 *ori*, T7 promoter, MCS1 and MCS2, $Amp^R$ | Novagen |
| pET28a | Expression vector, pBR322 *ori*, T7 promoter, MCS, $Kan^R$ | Novagen |
| pET28a-His-*hcp1*$^{PP}$ | *hcp1*$^{PP}$ encoding $Hcp1^{PP}$ with an N-terminal $His_6$ tag cloned into pET28a, $Kan^R$ | This study |
| pETDuet-1-His-*tssA1*$^{PP}$ | *tssA1*$^{PP}$ encoding $TssA1^{PP}$ with an N-terminal $His_6$ tag cloned into pETDuet-1 MCS1, $Amp^R$ | This study |
| pETDuet-1-V5-*tssA1*$^{PP}$ | *tssA1*$^{PP}$ encoding $TssA1^{PP}$ with a N-terminal V5 tag cloned into pETDuet-1 MCS1, $Amp^R$ | This study |
| pETDuet-1-*tagB1*$^{PP}$-StrepII | *tagB1*$^{PP}$ encoding $TagB1^{PP}$ with a C-terminal StrepII tag cloned into pETDuet-1 MCS2, $Amp^R$ | This study |
| pETDuet-1-*tagB1*$^{PP}$-V5 | *tagB1*$^{PP}$ encoding $TagB1^{PP}$ with a C-terminal V5 tag cloned into pETDuet-1 MCS2, $Amp^R$ | This study |
| pETDuet-1-His-*tssA1*$^{PA}$ | *tssA1*$^{PA}$ encoding $TssA1^{PA}$ with an N-terminal $His_6$ tag cloned into pETDuet-1 MCS1, $Amp^R$ | This study |
| pETDuet-1-V5-*tssA1*$^{PA}$ | *tssA1*$^{PA}$ encoding $TssA1^{PA}$ with an N-terminal V5 tag cloned into pETDuet-1 MCS1, $Amp^R$ | This study |
| pETDuet-1-*tagJ1*$^{PA}$-StrepII | *tagJ1*$^{PA}$ encoding $TagJ1^{PA}$ with a C-terminal StrepII tag cloned into pETDuet-1 MCS2, $Amp^R$ | This study |
| pETDuet-1-*tagJ1*$^{PA}$-V5 | *tagJ1*$^{PA}$ encoding $TagJ1^{PA}$ with a C-terminal V5 tag cloned into pETDuet-1 MCS2, $Amp^R$ | This study |
| pCcmG1 | L36-A184 of *Escherichia coli* CcmG expressed from pET22b(+), $Amp^R$ | Mavridou lab |
| mini-CTX1-P*lac* | Plasmid for the integration of genes into the *att* site of pseudomonal chromosomes, pMB1-derived *ori*, P*lac*, $Tet^R$ | (5) |
| mini-CTX1-P*lac*-*tssB1*$^{PP}$-*mScarlet-I* | *tssB1*$^{PP}$ encoding $TssB1^{PP}$ C-terminally fused to m-Scarlet-I cloned into miniCTX-1-P*lac*, $Tet^R$ | This study |
| pRL662-*gfp2* | pRL662 (35) derivative expressing GFP2, $Gent^R$ | Erh-Min Lai lab |
| pKNG101 | Gene replacement suicide vector, R6K *ori*, *sacB*, $Str^R$ | (4) |
| pUT18 | Bacterial-two-hybrid vector encoding the T18 fragment of *Bordetella pertussis* adenylate cyclase to construct N-terminal fusions, $Amp^R$ | (13) |
| pUT18C | Bacterial-two-hybrid vector encoding the T18 fragment of *Bordetella pertussis* adenylate cyclase to construct C-terminal fusions, $Amp^R$ | (13) |
| pKNT25 | Bacterial-two-hybrid vector encoding the T25 fragment of *Bordetella pertussis* adenylate cyclase to construct N-terminal fusions, $Kan^R$ | (13) |
| pUT18C_$TagB1^{PP}$ | T18 fragment of *Bordetella pertussis* adenylate cyclase C-terminally fused to $TagB1^{PP}$, $Amp^R$ | This study |
| pUT18_$TagB1^{PP}$ | T18 fragment of *Bordetella pertussis* adenylate cyclase N-terminally fused to $TagB1^{PP}$, $Amp^R$ | This study |

| | | |
|---|---|---|
| pKNT25_TssE1$^{PP}$ | T25 fragment of *Bordetella pertussis* adenylate cyclase N-terminally fused to TssE1$^{PP}$, Kan$^{R}$ | This study |
| pKNT25_TssF1$^{PP}$ | T25 fragment of *Bordetella pertussis* adenylate cyclase N-terminally fused to TssF1$^{PP}$, Kan$^{R}$ | This study |
| pKNT25_TssG1$^{PP}$ | T25 fragment of *Bordetella pertussis* adenylate cyclase N-terminally fused to TssG1$^{PP}$, Kan$^{R}$ | This study |
| pKNT25_TssK1$^{PP}$ | T25 fragment of *Bordetella pertussis* adenylate cyclase N-terminally fused to TssK1$^{PP}$, Kan$^{R}$ | This study |
| pKNT25_Hcp1$^{PP}$ | T25 fragment of *Bordetella pertussis* adenylate cyclase N-terminally fused to Hcp1$^{PP}$, Kan$^{R}$ | This study |
| pKNT25_VgrG1$^{PP}$ | T25 fragment of *Bordetella pertussis* adenylate cyclase N-terminally fused to VgrG1$^{PP}$, Kan$^{R}$ | This study |
| pKNT25_TssA1$^{PP}$ | T25 fragment of *Bordetella pertussis* adenylate cyclase N-terminally fused to TssA1$^{PP}$, Kan$^{R}$ | This study |
| pKNG101-*tssA1*$^{PP}$ | PCR fragment containing the regions upstream and downstream *tssA1*$^{PP}$ cloned in pKNG101 for double recombination; when inserted into the chromosome and the plasmid cured, the strain is a *tssA1*$^{PP}$ mutant, Str$^{R}$ | (22) |
| pKNG101-*tagB1*$^{PP}$ | PCR fragment containing the regions upstream and downstream *tagB1*$^{PP}$ cloned in pKNG101; when inserted into the chromosome and the plasmid cured, the strain is a *tagB1*$^{PP}$ mutant, Str$^{R}$ | This study |
| pKNG101-*hcp1*$^{PP}$ | PCR fragment containing the regions upstream and downstream *hcp1*$^{PP}$ cloned in pKNG101; when inserted into the chromosome and the plasmid cured, the strain is a *hcp1*$^{PP}$ mutant, Str$^{R}$ | This study |
| pKNG101-*tssB1*$^{PP}$*::sfgfp* | PCR fragment containing the 3´ region of *tssB1*$^{PP}$ fused to *sfgfp* and the downstream region of the same gene cloned in pKNG101; when inserted into the chromosome and the plasmid cured, the strain expresses TssB1$^{PP}$ C-terminally fused to sfGFP, Str$^{R}$ | This study |
| pKNG101-*tssB1*$^{PP}$*::mScarlet-I* | PCR fragment containing the 3´ region of *tssB1*$^{PP}$ fused to *mScarlet-I* and the downstream region of the same gene cloned in pKNG101, when inserted into the chromosome and the plasmid cured, the strain expresses TssB1$^{PP}$ C-terminally fused to mScarlet-I, Str$^{R}$ | This study |
| pKNG101-*tssA1*$^{PP}$*::sfgfp* | PCR fragment containing the 5´ region of *tssA1*$^{PP}$ fused to *sfgfp* and the upstream region of the same gene cloned in pKNG101, when inserted into the chromosome and the plasmid cured, the strain expresses TssA1$^{PP}$ N-terminally fused to sfGFP, Str$^{R}$ | This study |
| pKNG101-*tagB1*$^{PP}$*::sfgfp* | PCR fragment containing the 3´ region of *tagB1*$^{PP}$ fused to *sfgfp* and the downstream region of the same gene cloned in pKNG101, when inserted into the chromosome and the plasmid cured, the strain expresses TagB1$^{PP}$ C-terminally fused to sfGFP, Str$^{R}$ | This study |
| pKNG101-*tssB1*$^{PA}$*::sfgfp* | PCR fragment containing the 3´ region of *tssB1*$^{PA}$ fused to *sfgfp* and the downstream region of the same gene cloned in pKNG101, when inserted into the chromosome and the plasmid cured, the strain | This study |

| | | |
|---|---|---|
| | expresses $TssB1^{PA}$ C-terminally fused to sfGFP, $Str^R$ | |
| pKNG101-*tssB2*$^{PA}$*::sfgfp* | PCR fragment containing the 3´ region of *tssB2*$^{PA}$ fused to *sfgfp* and the downstream region of the same gene cloned in pKNG101, when inserted into the chromosome and the plasmid cured, the strain expresses $TssB2^{PA}$ C-terminally fused to sfGFP, $Str^R$ | This study |
| pKNG101-*tssB3*$^{PA}$*::sfgfp* | PCR fragment containing the 3´ region of *tssB3*$^{PA}$ fused to *sfgfp* and the downstream region of the same gene cloned in pKNG101, when inserted into the chromosome and the plasmid cured, the strain expresses $TssB3^{PA}$ C-terminally fused to sfGFP, $Str^R$ | This study |
| pKNG101-*tagJ1*$^{PA}$*::sfgfp* | PCR fragment containing the 3´ region of *tagJ1*$^{PA}$ fused to *sfgfp* and the downstream region of the same gene cloned in pKNG101, when inserted into the chromosome and the plasmid cured, the strain expresses $TagJ1^{PA}$ C-terminally fused to sfGFP, $Str^R$ | This study |
| pKNG101-*tagB1*$^{PP}$*::(StrepII)*$_2$ | PCR fragment containing the 3´ region of *tagB1*$^{PP}$ fused to a twin StrepII tag and the downstream region of the same gene cloned in pKNG101, when inserted into the chromosome and the plasmid cured, the strain expresses $TagB1^{PP}$ C-terminally fused to two consecutives StrepII tags, $Str^R$ | This study |
| pRK600 | Helper plasmid, ColE1 *ori*, *mob*RK2, *tra*RK2, $Cam^R$ | (36) |

**Table S3.** Oligonucleotide primers used in this study. The "Brief description" column provides basic information on the primer design (restriction enzyme(s) used for cloning, encoded protein or gene replaced by antibiotic resistance cassette, forward or reverse orientation of the primer (F or R)). Primers marked with TE (for "tag exchange") were used for replacing an existing affinity tag with an alternative one, primers marked with UP, DOWN or MED were used for construction of mutant strains, primers marked with T18 or T25 were used for construction of plasmids used in bacterial-two-hybrid experiments, and primers marked with CTX were used for the construction of miniCTX vector derivatives for insertion of genes into the *Pseudomonas* genome.

| Number | Brief description | Sequence (5´-3´) |
|---|---|---|
| P1 | NdeI.Hcp1$^{PP}$.F | ggccggcatatgatgttgttaatggagagtttc |
| P2 | BamHI.Hcp1$^{PP}$.R | tacgcaggatccggttcggtacgtgcttag |
| P3 | NdeI.TagB1$^{PP}$.F | aattaacatatggacgatgcaccaatgac |
| P4 | XhoI.TagB1$^{PP}$StrepII.R | ttaattctcgagctacttttcgaactgcgggtggctccacagtccttcgatcgtcagc |
| P5 | EcoRI.TssA1$^{PP}$.F | gcgctgaattcgcctgaaattctcaatgacctc |
| P6 | HindIII.TssA1$^{PP}$.R | tggttaagcttgacgttcgtctcaaggctt |
| P7 | NdeI.TagJ1$^{PA}$.F | gcttacatatggccgacccttcattcgc |
| P8 | XbaI.TagJ1$^{PA}$StrepII.R | aaaaaagatctacttttcgaactgcgggtggctccaggcgtccgtcggctcgaag |
| P9 | EcoRI.TssA1$^{PA}$.F | atcgagaattccgtgctggatgtacccg |
| P10 | HindIII.TssA1$^{PA}$.R | tggccaagcttccgctaggcgttactcg |
| P11 | TE.TssA1$^{PP}$V5.F | accgaggagagggttagggataggcttacccatggtatatctccttcttaaagttaaac |
| P12 | TE.TssA1$^{PP}$V5.R | cctaaccctctcctcggtctcgattctacgcctgaaattctcaatgacctctcgctg |
| P13 | TE.TagB1$^{PP}$V5.F | accgaggagagggttagggataggcttacccagtccttcgatcgtcagcagcaggcgctg |
| P14 | TE.TagB1$^{PP}$V5.R | cctaaccctctcctcggtctcgattctacgtagctcgagtctggtaaagaaaccgctgctgc |
| P15 | TE.TssA1$^{PA}$V5.F | accgaggagagggttagggataggcttacccatggtatatctccttcttaaagttaaac |
| P16 | TE.TssA1$^{PA}$V5.R | cctaaccctctcctcggtctcgattctacgctggatgtacccgttttgctggctg |
| P17 | TE.TagJ1$^{PA}$V5.F | accgaggagagggttagggataggcttaccggcgtccgtcggctcgaagtccagttcac |
| P18 | TE.TagJ1$^{PA}$V5.R | cctaaccctctcctcggtctcgattctacgtagatctcaattggatatcggccggccacgcg |
| P19 | XbaI.tagB1$^{PP}$UP.F | ggttaatctagagctgcgccggtgtt |
| P20 | tagB1$^{PP}$UP.R | ttacagtccatcgtccatgtgcctgct |
| P21 | tagB1$^{PP}$DOWN.F | atggacgatggactgtaacgcgcagcg |
| P22 | BamHI.tagB1$^{PP}$DOWN.R | ttatcaggatccgtcgccggcctgtt |
| P23 | XbaI.tagB1$^{PP}$-gfpUP.F | ggttcctctagactggtcgaactctacgaacgt |
| P24 | tagB1$^{PP}$-gfpUP.R | tgctgctgccagtccttcgatcgtcag |
| P25 | tagB1$^{PP}$-gfpMED.F | gaaggactggcagcagcaggaggagga |
| P26 | tagB1$^{PP}$-gfpMED.R | cgctgcgcgtcattacttatacagctc |
| P27 | tagB1$^{PP}$-gfpDOWN.F | ataagtaatgacgcgcagcgcccggcacc |
| P28 | BamHI.tagB1$^{PP}$-gfpDOWN.R | aattaaggatccttcacggccgaattccttgc |
| P29 | XbaI.tssA1$^{PP}$-gfpUP.F | ggttcctctagaacaaggactggatcgaagtgc |
| P30 | tssA1$^{PP}$-gfpUP.R | tttacgcatagaacactctggacatgg |

| | | |
|---|---|---|
| P31 | tssA1$^{PP}$-gfpMED.F | gagtgttctatgcgtaaaggcgaagaa |
| P32 | tssA1$^{PP}$-gfpMED.R | tcctcctcctgctgctgccttatacagctcgtccat |
| P33 | tssA1$^{PP}$-gfpDOWN.F | gcagcagcaggaggaggacctgaaattctcaatgac |
| P34 | BamHI.tssA1$^{PP}$-gfpDOWN.R | aattaaggatcctgctgtctgcatcgccttg |
| P35 | XbaI.tssB1$^{PP}$-gfpUP.F | ggttcctctagattcgcagcgaattgctcg |
| P36 | tssB1$^{PP}$-gfpUP.R | tgctgctgcggcgcgatcgtcgctgga |
| P37 | tssB1$^{PP}$-gfpMED.F | gatcgcgccgcagcagcaggaggagga |
| P38 | tssB1$^{PP}$-gfpMED.R | ggctcctgcttacttgtacagctcgtc |
| P39 | tssB1$^{PP}$-gfpDOWN.F | tacaagtaagcaggagcccgaacatgac |
| P40 | BamHI.tssB1$^{PP}$-gfpDOWN.R | aattaaggatccagtcaccgatcaacatgccg |
| P41 | tssB1$^{PP}$-mScMED.F | gatcgcgccgcagcagcaggaggaggaatggttagcaaagg cgaggc |
| P42 | tssB1$^{PP}$-mScMED.R | ggctcctgcttaggatcccttatacagttcgtccatg |
| P43 | tssB1$^{PP}$-mScDOWN.F | ggatcctaagcaggagcccgaacatgaccgacaagc |
| P44 | SacI.tssB1$^{PP}$-mScCTX.F | aattatcccgggtaacaggaggaattaaccatgattcgcagcga attgctcg |
| P45 | XmaI.tssB1$^{PP}$-mScCTX.R | aattccgagctctgcttaggatcccttatacagttcgtc |
| P46 | XbaI.tssB2$^{PA}$-gfpUP.F | ggttcctctagattatcgcaggctgtgccacc |
| P47 | tssB2$^{PA}$-gfpUP.R | tgctgctgcgtcctgggagggggcggc |
| P48 | tssB2$^{PA}$-gfpMED.F | tcccaggacgcagcagcaggaggagga |
| P49 | tssB2$^{PA}$-gfpMED.R | ctagggggtggcttacttatacagctcgtccataccg |
| P50 | tssB2$^{PA}$-gfpDOWN.F | aagtaagccaccccctagccaaggaag |
| P51 | BamHI.tssB2$^{PA}$-gfpDOWN.R | aattaaggatcccgctggagacgtattgcatc |
| P52 | XbaI.tssB3$^{PA}$-gfpUP.F | ggttcctctagagggactgcgatgatcagggcatcc |
| P53 | tssB3$^{PA}$-gfpUP.R | tgctgctgcggccggctggtcggccgg |
| P54 | tssB3$^{PA}$-gfpMED.F | cagccggccgcagcagcaggaggagga |
| P55 | tssB3$^{PA}$-gfpMED.R | ccgggaagagggttacttatacagctcgtccataccg |
| P56 | tssB3$^{PA}$-gfpDOWN.F | aagtaaccctcttcccggagaagccgc |
| P57 | BamHI.tssB3$^{PA}$-gfpDOWN.R | aattaaggatccagcaggccgatgtcctgc |
| P58 | XbaI.tagJ1$^{PA}$-gfpUP.F | aaaaatctagagccgatggtacagacctactccaccg |
| P59 | tagJ1$^{PA}$-gfpUP.R | tgctgctgcggcgtccgtcggctcgaagtcc |
| P60 | tagJ1$^{PA}$-gfpMED.F | ccgacggacgccgcagcagcaggaggagga |
| P61 | tagJ1$^{PA}$-gfpMED.R | cgcaggtcgaacttacttatacagctcgtccataccg |
| P62 | tagJ1$^{PA}$-gfpDOWN.F | tataagtaagttcgacctgcgtgaactggacttcgagc |
| P63 | BamHI.tagJ1$^{PA}$-gfpDOWN.R | tataaggatccgccggtctccaggtccaggttg |
| P64 | XbaI.T18-tagB1.F | gcggtctagaggacgatgcaccaatgacaca |
| P65 | BamHI.T18-tagB1.R | gcgtggatccttacagtccttcgatcgtcag |
| P66 | XbaI.tagB1-T18.F | gcgatctagagatggacgatgcaccaatgac |
| P67 | BamHI.tagB1-T18.R | atatggatcctccagtccttcgatcgtcag |
| P68 | XbaI.tssE1-T25.F | gcgcgtctagagagtaaccaggcccgcctgct |
| P69 | BamHI.tssE1-T25.R | atatggatcctcgttgaggttctgcactttg |
| P70 | XbaI.tssF1-T25.F | gcgcgtctagagctcgacgaactgctgcccta |
| P71 | BamHI.tssF1-T25.R | atatggatcctcgaccaggggctgatc |
| P72 | XbaI.tssG1-T25.F | gcgcgtctagaggccagcacgcaccggcgatca |
| P73 | BamHI.tssG1-T25.R | atatggatcctccaaggcgaccccatcgtaag |
| P74 | XbaI.tssK1-T25.F | gcgcgtctagagagcaagcagagccgggtgat |
| P75 | BamHI.tssK1-T25.R | atatggatcctctttgagcaccgccatcagtt |
| P76 | XbaI.hcp1-T25.F | cgcgctctagagttgttaatggagagtttcacaatg |
| P77 | BamHI.hcp1-T25.R | atatggatcctcggcaaacactttgttcgcgg |
| P78 | XbaI.vgrG1-T25.F | gcgcgtctagagctaactgacgttctttcc |
| P79 | BamHI.vgrG1-T25.R | atatggatcctcgtcgttgatccgcactgc |
| P80 | XbaI.tssA1-T25.F | gcgcgtctagagcctgaaattctcaatgacc |

| | | |
|---|---|---|
| P81 | BamHI.tssA1-T25.R | atatggatcctcggtgcccagcacatcgcg |
| P82 | XbaI.tssB1$^{PA}$-gfpUP.F | gggaagcactaccagcagtcagaagttcat |
| P83 | tssB1$^{PA}$-gfpUP.R | cctgctgctgccgcctgcgg |
| P84 | tssB1$^{PA}$-gfpMED.F | gcagcagcaggaggaggaatgcgtaaaggcgaagaa |
| P85 | tssB1$^{PA}$-gfpMED.R | atcctctcattacttatacagctcgtccat |
| P86 | tssB1$^{PA}$-gfpDOWN.F | taatgagaggattccagcatggccgaattgag |
| P87 | BamHI.tssB1$^{PA}$-gfpDOWN.R | gtagtagtcgccgaccaggcagcc |
| P88 | XbaI.tagB1$^{PP}$-StrepII2UP.F | ggttcctctagattggcgaatacctgcaaggc |
| P89 | tagB1$^{PP}$-StrepII2UP.R | gtggctccacttttcgaactgcgggtggctccacagtccttcgat cgtcagca |
| P90 | tagB1$^{PP}$-StrepII2DOWN.F | ttcgaaaagtggagccacccgcagttcgaaaagtagctgacga tcgaaggactg |
| P91 | BamHItagB1$^{PP}$-StrepII2DOWN.R | aattaaggatcctgcggtttgcttgtcggtc |
| P92 | XbaI.hcp1$^{PP}$UP.F | taatctagactgcgcgaccgctac |
| P93 | hcp1$^{PP}$UP.R | gcgcggaattcgtaaagggttgctcatgc |
| P94 | hcp1$^{PP}$DOWN.F | gcagaattcgcacgtaccgaaccat |
| P95 | BamHIhcp1$^{PP}$DOWN.R | aatggatccttgcggttcgctctg |

### LEGENDS FOR SUPPLEMENTARY VIDEOS

**Video S1. *In vivo* imaging of *P. putida rpoN* TssB1-sfGFP.** The full cycle of T6SS assembly, including sheath initiation, extension, contraction and disassembly over approximately 120 seconds is shown for a representative sheath. This video was generated using a fluorescence microscopy time-lapse recording (total time 140 seconds) of *P. putida rpoN* expressing TssB1-sfGFP from the native *tssB1* locus. Images were acquired every 2 s and the video is presented at 7 frames/s.

**Video S2. *In vivo* imaging of *P. putida rpoN* TagB1-sfGFP.** TagB1 forms transient foci which accumulate and abruptly disappear over time. Disappearance of foci is not due to photobleaching as new TagB1-sfGFP foci appear in the same or neighboring cells. This video was generated using a fluorescence microscopy time-lapse recording (total time 120 seconds) of *P. putida rpoN* expressing TagB1-sfGFP from the native *tagB1* locus. Images were acquired every 2 s and the video is presented at 7 frames/s.

### LEGENDS FOR SUPPLEMENTARY FILES

**File S1. Overview of the results originating from an *in silico* analysis of the structural components of 100 T6SS clusters.** The T6SS clusters included in the analysis are organized by phylogenetic group (groups 1-5 are shown in different tabs, with phylogenetic subgroups 4A and 4B shown separately). 20 T6SS clusters per phylogenetic group were analyzed and a hyperlink to the SecReT6 website (19) is given for each cluster; strains appearing in multiple phylogenetic groups have multiple T6SSs. The T6SS cluster that is best characterized in each phylogenetic group (highlighted in grey) was set as the reference cluster for the analysis (distinct clusters from *P. aeruginosa*, *E. coli*, *P. putida* and *A. tumefaciens* were selected as references). White cells indicate proteins that cannot be found in a given T6SS cluster. In some cases, more than one analogue of a specific T6SS core component is found in a given cluster; for example there are two TssC analogues in all T6SS clusters of phylogenetic group 5, several T6SS clusters have multiple VgrG or Hcp components, and in approximately 2% of cases there are two analogues of some components of the membrane complex or the baseplate. These additional components are not shown for simplicity and the protein that is most similar to that of the reference genome is included in the spreadsheet.

**File S2. *P. putida* TagB1-(StrepII)$_2$ specifically co-purifies with TssA1.** Proteins eluted from Strep-Tactin Sepharose resin incubated with *P. putida rpoN* or *P. putida rpoN tagB1-(StrepII)$_2$* whole-cell lysates, as identified and quantified by mass spectrometry. The file contains the output of a volcano plot analysis, performed in Perseus version 1.5.8.5 (9); proteins identified in the TagB1-(StrepII)$_2$ eluate were compared to those identified in the control eluate (untagged *P. putida* TagB1). Proteins with a positive "difference" value are enriched in the TagB1-(StrepII)$_2$ condition compared to the control, whilst proteins with a negative "difference" value are enriched in the control compared to the TagB1-(StrepII)$_2$ condition. Proteins with significantly different abundances in the two conditions are marked "YES" in the "Significant" column. Log$_2$ LFQ intensity values, in technical triplicate, are given for each protein sample in each condition.
